# Dynamic localization of SPO11-1 and conformational changes of meiotic axial elements during recombination initiation of maize meiosis

**DOI:** 10.1101/489278

**Authors:** Jia-Chi Ku, Arnaud Ronceret, Inna Golubovskaya, Ding Hua Lee, Chiting Wang, Ljudmilla Timofejeva, Yu-Hsin Kao, Ana Karen Gomez Angoa, Karl Kremling, Rosalind Williams-Carrier, Robert Meeley, Alice Barkan, W. Zacheus Cande, Chung-Ju Rachel Wang

**Affiliations:** Institute of Plant and Microbial Biology, Academia Sinica, Taipei 11529, Taiwan; Department of Molecular and Cell Biology and Plant and Microbial Biology, University of California, Berkeley, CA 94720, USA; Instituto de Biotecnología / UNAM Cuernavaca, Morelos 62210 Mexico; N.I. Vavilov Institute of Plant Industry, St. Petersburg 190000, Russia; Institute of Molecular Biology, University of Oregon, Eugene, OR 97403, USA; Corteva Agriscience, Johnston, Iowa 50131, USA

**Keywords:** Maize, SPO11, recombination, axial element, meiosis

## Abstract

Meiotic double-strand breaks (DSBs) are generated by the evolutionarily conserved SPO11 complex in the context of chromatin loops that are organized along axial elements (AEs) of chromosomes. However, how DSBs are formed with respect to chromosome axes and the SPO11 complex remains unclear in plants. Here, we confirm that DSB and bivalent formation are defective in maize *spo11-1* mutants. Super-resolution microscopy demonstrates dynamic localization of SPO11-1 during recombination initiation, with variable numbers of SPO11-1 foci being distributed in nuclei but similar numbers of SPO11-1 foci being found on AEs. Notably, cytological analysis of *spo11-1* meiocytes revealed an aberrant AE structure. At leptotene, AEs of wild-type and *spo11-1* meiocytes were similarly curly and discontinuous. However, during early zygotene, wild-type AEs become uniform and exhibit shortened axes, whereas the elongated and curly AEs persisted in *spo11-1* mutants, suggesting that loss of SPO11-1 compromised AE structural maturation. Our results reveal an interesting relationship between SPO11-1 loading onto AEs and the conformational remodeling of AEs during recombination initiation.

**Author Summary:** Meiosis is essential during sexual reproduction to produce haploid gametes. Recombination is the most crucial step during meiotic prophase I. It enables pairing of homologous chromosomes prior to their reductional division and generates new combinations of genetic alleles for transmission to the next generation. Meiotic recombination is initiated by generating DNA double-strand breaks (DSBs) via SPO11, a topoisomerase-related enzyme. The activity, timing and location of this DSB machinery must be controlled precisely, but how this is achieved remains obscure. Here, we show dynamic localization of SPO11-1 on chromatin during meiotic initiation in maize, yet a similar number of SPO11-1 is able to load onto axial elements (AEs), which accompanies a structural change of the AEs of wild-type meiotic chromosomes. Interestingly, loss of SPO11-1 not only affects DSB formation but also impairs structural alterations of AEs, resulting in abnormally long and curly AEs during early meiosis. Our study provides new insights into SPO11-1 localization during recombination initiation and suggests an intimate relationship between DSB formation and AE structural changes.

## Introduction

Meiosis is an essential cell division that halves ploidy numbers through two successive chromosome divisions to produce gametes for sexual reproduction. The faithful segregation of homologous chromosomes requires meiotic recombination during prophase I, which mediates pairing of homologous chromosomes and creates physical connections until their segregation [1]. Meiotic recombination is initiated by the introduction of double-strand breaks (DSBs) into chromosomal DNA via SPO11, a protein of the topoisomerase VI family [2]. The SPO11 complex cuts double-stranded DNA and links covalently to the 5’ ends at the break point [3]. Following SPO11 removal together with 5’-oligonucleotides, DSBs are further resected to produce 3’ single-stranded DNA (ssDNA) tails for searching homologues with the assistance of RAD51 and DMC1 recombinase proteins. Only a small subset of DSBs are repaired using a chromatid of homologous chromosomes to give rise to crossovers (COs), whereas most DSBs lead to non-crossovers (NCOs) [4]. These CO sites, seen as chiasmata at the diplotene stage, ensure that pairs of homologs, i.e. the bivalents, maintain their association at the metaphase plate, which allows subsequent reductional segregation at anaphase I. Thus, at least one CO per bivalent is needed for accurate segregation of homologous chromosomes. When DSB formation is defective, such as in *spo11* mutants, meiotic recombination is impaired in all species analyzed to date, leading to infertile or aneuploid gametes [5–10].

These essential meiotic DSBs are also potentially harmful to cells, so their formation is tightly regulated both temporally and spatially to allow proper initiation of recombination [1]. For example, the timing of DSB induction is linked to the completion of pre-meiotic DNA replication [11] and it is restricted to a narrow timeframe [12]. Moreover, the total number of DSBs is also regulated, as too few DSBs could be detrimental given that sufficient DSBs are required for homologous engagement in most species. However, too many DSBs can also be deleterious since they may result in chromosome fragmentation if unrepaired. Furthermore, clustered DSBs in a short interval is also likely prevented [13]. Recent studies in yeast and mouse have revealed that DNA damage-responsive kinase Tel1/ATM activity and subsequent homologous engagement are involved in negative feedback loops to restrain SPO11 from marking too many DSBs [13–15]. However, the control mechanism and its regulatory network for initiating and then halting DSB formation are not yet well understood.

During meiotic prophase in most eukaryotes studied, axial elements (AEs) are assembled soon after the pre-meiotic S phase to organize sister chromatids into arrays of chromatin loops [16, 17]. Although it remains unclear how the DSB machinery is regulated, several yeast chromosomal axis-located proteins such as Rec114, Mer2, Red1, Hop1 and Mei4 have been found to be required for DSB formation [18]. These findings suggest that the DSB machinery may be associated with chromosome axes and that this association appears to be necessary for SPO11 transesterase activity. Interestingly, DNA sequences prone to DSBs (i.e. DSB hotspots) have been mapped to loop domains away from the axis-associated sequences [19]. Thus, a ‘tethered loop-axis complex’ model has been postulated in an attempt to reconcile these findings, which argues for a transient but as yet unobserved localization of the SPO11 complex on AEs during DSB formation [20, 21]. The finding that yeast Spp1 protein interacts with AE-located Mer2 and also recognizes H3K4 methylation marks which are enriched around DSB hotspots provides a molecular model of how tethering is achieved [22, 23].

The meiotic chromosome axis consists of a cohesion complex that holds sister chromatids and other axial proteins that organize chromosomes into separate units [24, 25]. In plants, the meiotic cohesin subunit REC8 α-kleisin has been shown to be required for AE formation [26, 27]. Several non-cohesin axial proteins that are assembled along with meiotic cohesins have also been isolated in plants. AtASY1/OsPAIR2 containing the HORMA (<u>Ho</u>p1, <u>R</u>ev7, <u>Ma</u>d2) domain was found to be important for CO formation [28, 29], and it is thought to be a homolog of yeast Hop1 [30]. Notably, plant ASY1/PAIR2 and yeast HOP1 contain conserved S/TQ motifs that can be phosphorylated by ATM/ATR kinase upon DNA damage [31]. Like other HORMA proteins, AtASY1/OsPAIR2 preferentially associates with unsynapsed chromosome axes and its protein level diminishes after synapsis [29, 32, 33]. Other axial proteins, including OsPAIR3/AtASY3/ZmDSY2 and AtASY4, contain coiled-coil domains and are required for proper axis formation and recombination, perhaps representing functional homologs of budding yeast Red1 and mammalian SYCP2/3 [34–39]. As meiotic prophase I proceeds, the axes of homolog are gradually brought into close juxtaposition during homologous pairing and then are tightly linked by synaptonemal complex (SC) which consists of two lateral elements (called axial elements prior to synapsis) separated by a central region where transverse filaments such as the ZIP1/ZYP1 protein are closely packed [17, 25].

The ‘tethered loop-axis complex’ model explains that DSBs are made upon chromatin loops being tethered with chromosome axes [40], but supporting cytological evidence is scarce. It remains unclear how DSBs are generated with respect to chromosome axes and SPO11 complex. In addition, how DSB sites communicate within a nucleus in order to avoid excess or clustered DSBs remains poorly understood. If tethering DNAs to chromosome axes is indispensable for DSB formation, how do the axes respond, if at all, to DSB formation? In species such as maize, barley and wheat with large chromosomes, subtle changes of chromosome structure and AE morphology are more easily observable than in species having small chromosomes [41–46]. Here, we show that, as expected, maize *Spo11-1* is required for DSB and bivalent formation. Interestingly, cytological analysis of *spo11-1* mutants revealed an unexpected phenotype in that AEs were longer than wild-type and had a curly appearance before promiscuous synapsis. Super-resolution microscopy showed that varying numbers of copious SPO11-1 foci are distributed in nuclei, but a relatively similar number of SPO11-1 foci were associated with AEs during the leptotene to zygotene transition. To our knowledge, these findings provide the first cytological evidence of dynamic SPO11-1 localization during DSB formation and suggest that a possible SPO11-1-dependent mechanism may mediate conformational changes of meiotic chromosome axes.

## Results

### Maize mutants *mtm99-14* and *mtm00-03* exhibit meiotic phenotypes

We used a forward genetic approach with active *Mutator* (*Mu*) transposition [47, 48] and identified *mtm99-14* and *mtm00-03* which are defective in male and female fertility (Tables S1 and S2). In maize, male meiosis occurs within anthers that develop progressively with respect to their positions on a male tassel. Meiotic progression, especially for prophase I, is correlated nicely with anther length, providing a reliable means of staging and comparing wild-type (WT) and mutant anthers [49]. The figure 1A displays that DAPI-stained chromosomes of leptotene meiocytes appear as long, thin threads with multiple spheres of heterochromatic knobs, and that a nucleolus is located within the nuclear interior. A profound chromosomal reorganization takes place in maize meiocytes at late leptotene, seen as that chromatin mass is polarized against an offset nucleolus and heterochromatic knobs exhibit transient elongation [50]. During WT zygotene, homologous chromosomes pair gradually, often manifesting as thicker chromosomes and cylinder-shaped heterochromatic knobs in synapsed regions. At pachytene, synapsis is complete with chromosomes exhibiting a uniform appearance along their lengths except for the heavily-stained rounded heterochromatic knobs. Meiosis in the *mtm00-03* and *mtm99-14* mutants appeared to progress normally until the zygotene stage, with an obviously abnormal chromosome phenotype in both mutants at pachytene stage (Fig 1A).

**Figure 1.**
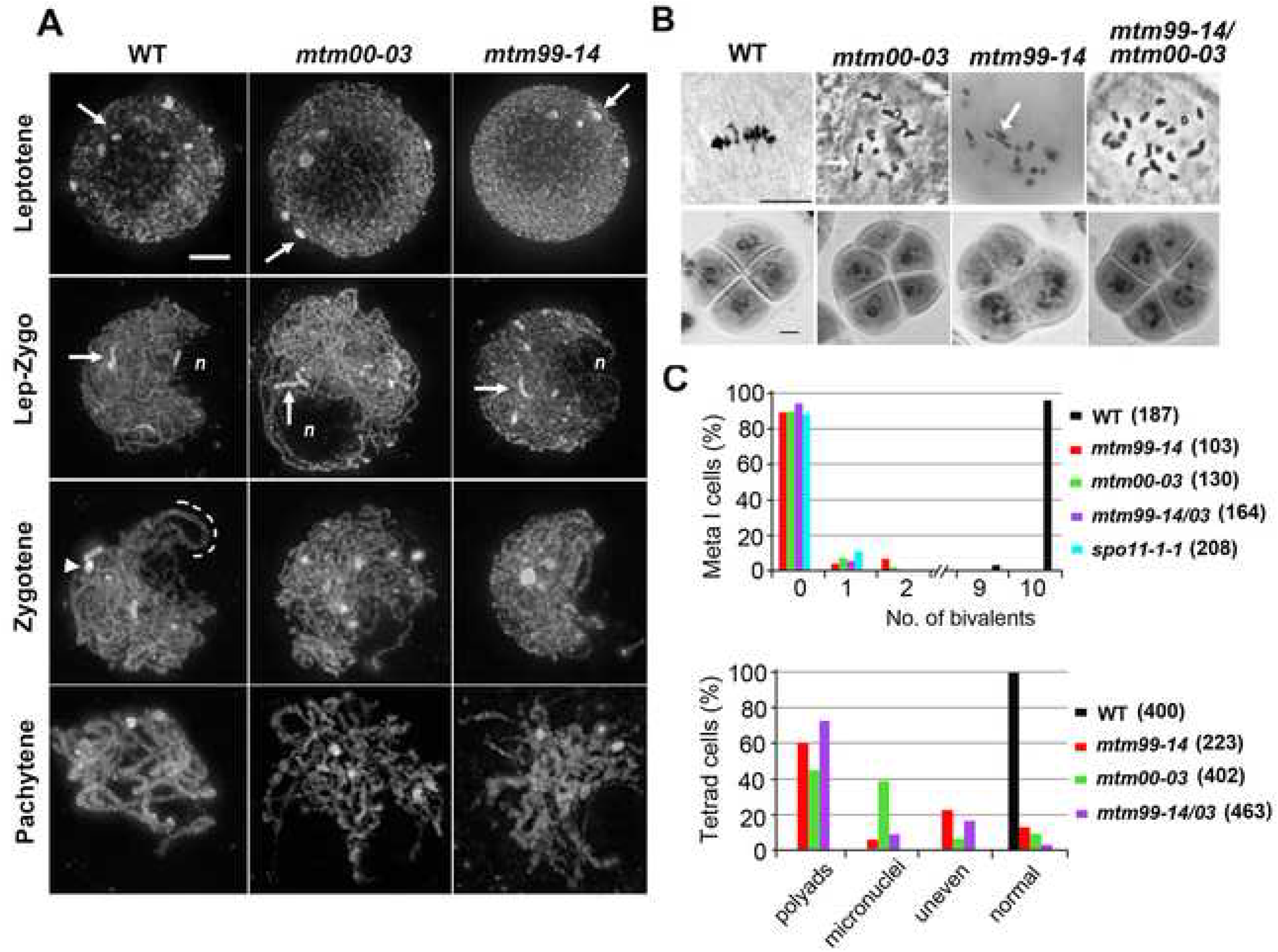
*mtm99-14* and *mtm00-03* are allelic mutants with defective meiosis. **(A)** Projected images of WT and mutant meiocytes stained with DAPI during early prophase I. Initially spherical heterochromatic knobs (arrows) become elongated at the leptotene-zygotene transition that accompanies a positional change of the nucleolus (*n*) from the nuclear interior to the periphery. At zygotene, paired knobs (arrowhead) and chromosomes (dashed line) were observed in WT. An obvious aberrant phenotype of both mutants was observed during the pachytene stage, manifested as fluffy chromosomes. Scale bar represents 5 μm. **(B)** Acetocarmine staining of male meiocytes during the metaphase I (upper panel) and tetrad stages (lower panel) of WT, *mtm00-03*, *mtm99-14*, and heteroallelic *mtm99-14/mtm00-03* plants. In WT male meiocytes, ten bivalents are aligned on a metaphase I plate and later produce normal tetrad cells. Mutant cells mainly present univalent chromosomes and rare interconnected chromosomes counted as bivalents (arrows). Scale bar represents 10 μm. **(C)** Quantitative results of meiotic phenotypes during the metaphase I and tetrad stages. Numbers of cells analyzed are shown in parentheses. The *spo11-1-1* is another mutant allele (see next section).

At the metaphase I, ten bivalents align on the metaphase plate in WT, but *mtm99-14* and *mtm00-03* meiocytes mainly contained univalents with occasional bivalents. In contrast to normal WT tetrad cells, mutants exhibited abnormal microspores as unequal tetrads, tetrads with micronuclei, or polyads (Fig 1B-1C). The reduced fertility of mutant maize ears suggested that female meiosis is also affected. Only a few seeds were obtained from mutants when they were pollinated with WT pollen (Table S2 and Fig S1). The proportion of seeds obtained from mutants is not statistically different to the expected proportion arising from completely random segregation of ten pairs of chromosomes (ε= 0.82 for *mtm99-14,* ε= 0.32 for *mtm00-03*, < ε_α5%_ =1.96). Given the similar phenotypes observed, we performed an allelism test and confirmed that the *mtm99-14* and *mtm00-03* mutants are allelic (Fig 1B, 1C and Tables S1, S2).

### Identification of mutations in the *Spo11-1* locus

We mapped the *mtm00-03* mutation to chromosome 5 by molecular markers (see Material and Methods). To identify the gene, *Mu* transposon insertions that are associated with the sterile phenotype were isolated using the *Mu*-Illumina approach [51]. We found that the *mtm99-14* and *mtm00-03* mutants harbor traces of *Mu* sequences associated with deletions spanning the *Spo11-1* (*GRMZM2G129913*) and *Phytochrome C2* (*PhyC2, GRMZM2G129889*) genes (Fig 2A). We could not determine the lengths of those deletions, and only one *Mu* border was recovered in our *Mu*-Illumina sequence analyses. Nevertheless, reverse transcription PCR experiments confirmed that neither the *mtm99-14* nor *mtm00-03* mutant expressed the *Spo11-1* gene in meiotic anthers (Fig 2B). Although the *PhyC2* gene is deleted, the closely paralogous gene *PhyC1* (*GRMZM2G057935*) that shares 94% identity with *PhyC2* is expressed in both mutants and it may be sufficient for phytochrome C function (Fig S2).

**Figure 2.**
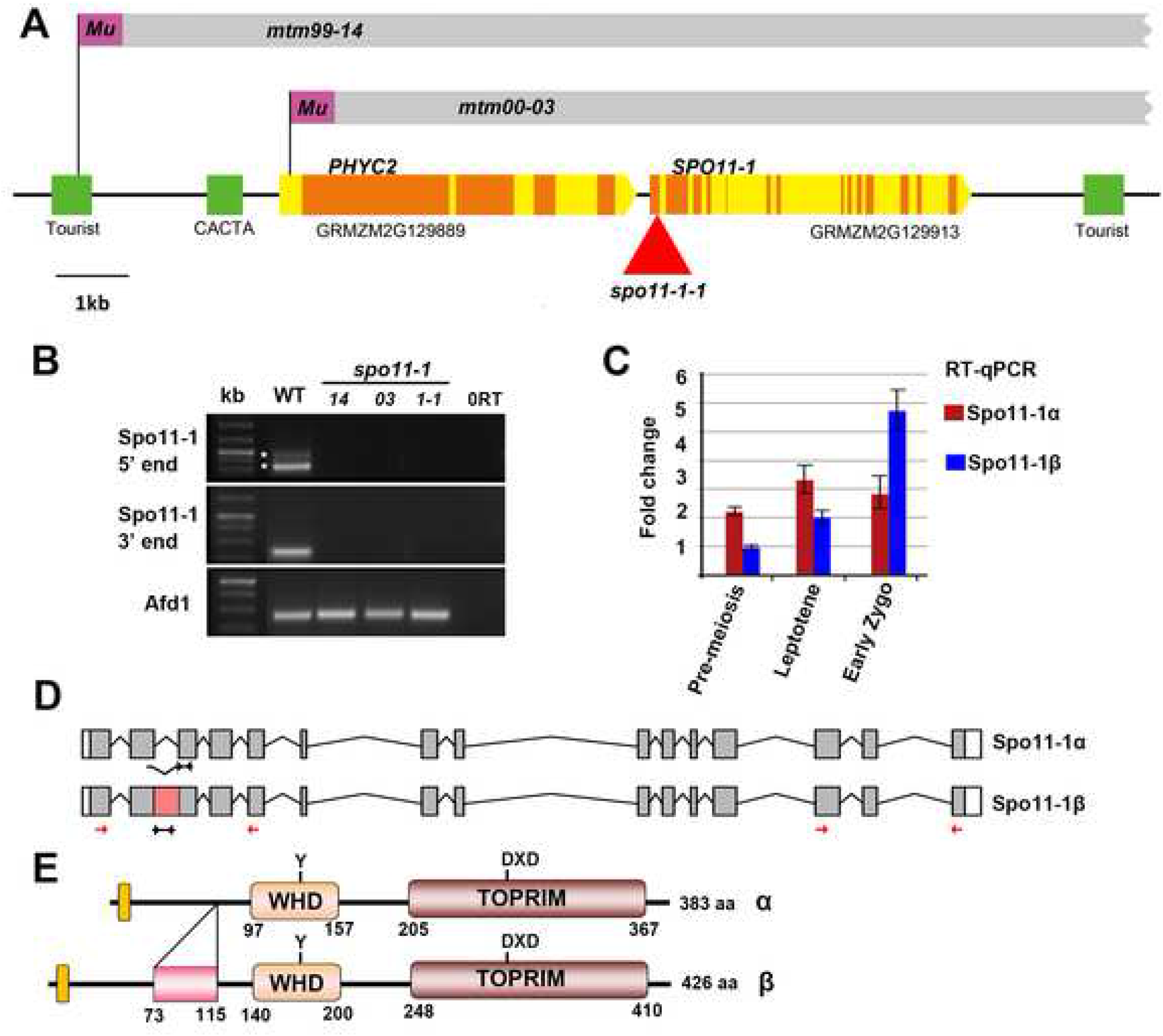
Structure and expression of the *SPO11-1* gene. **(A)** Schematic diagram of the maize genomic region. Both *mtm99-14* and *mtm00-03* mutants contain remnants of *Mu* sequences (purple boxes) that are associated with deletions (grey boxes) spanning both the *PhyC2* and *Spo11-1* genes. Coding regions and introns/UTRs are represented as orange and yellow boxes, respectively. The additional *Mu* insertion allele (*spo11-1-1*) is indicated by a red triangle. Green boxes represent transposable elements (Tourist and CACTA-like elements). **(B)** RT-PCR analyses to amplify the *Spo11-1* at 5’ and 3’ regions exhibited splicing variants (asterisks) in WT and confirmed that *Spo11-1* transcripts are absent in all mutant alleles. Positions of primer sets are indicated by red arrows in (D). *Afd1/Rec8* expression was used as a control. 0RT represents a negative control lacking cDNA. **(C)** Quantitative PCR using specific primers showed differential expression of the *Spo11-1α* and *Spo11-1β* transcripts. Positions of primer sets are indicated in by black arrows (D). **(D, E)** Gene models (D) and predicted proteins (E) of SPO11-1α and SPO11-1β are shown. The additional region of SPO11-1β (amino acid 73-115) is indicated by a pink box.

Considering complex rearrangements that may occur in both the *mtm99-14* and *mtm00-03* mutants, we searched for other mutations affecting only the *Spo11-1* gene. From the Trait Utility System for Corn (TUSC) transposon mutagenesis population [52], we identified one additional mutant, named *spo11-1-1,* containing a *Mu* insertion in the first exon (Fig 2A). We showed that the *spo11-1-1* mutant exhibited identical phenotypes to the *mtm99-14* and *mtm00-03* mutants, and an allelism test confirmed that the *spo11-1-1* mutant is allelic (Fig S3 and Table S1). Accordingly, we renamed *mtm99-14* and *mtm00-03* mutants as *spo11-1-Δ14* and *spo11-1-Δ03*, respectively. Since all alleles exhibited an identical phenotype, we characterized the *spo11-1-Δ03* mutant in a faster growing background extensively and included some results of other alleles in supplemental figures (Fig S3, S20 and S21).

### Maize *Spo11-1* exhibit alternative splicing

Our RT-PCR analysis of *Spo11-1* expression detected two spliced variants, which arose from retention of the second intron (Fig 2B, 2D). Both transcripts are predicted to produce functional SPO11-1 proteins because they contain a nuclear localization signal (KLRR), the topoisomerase-primase ‘TOPRIM’ domain that is essential for catalytic activity, and the conserved tyrosine residue in the winged-helix DNA-binding (WHD) domain (Fig 2E). The larger protein isoform (named Spo11-1β) contains an extra 43 amino-acid domain, which is predicted to form an alpha-helix motif upstream of the TopVIA type endonuclease domain (Fig S4, S5). We designed transcript-specific primers (Table S7) to examine temporal expression of the two transcript variants. Quantitative PCR results revealed that *Spo11*-*1α* is more strongly expressed during leptotene, whereas the *Spo11*-*1β* transcript level is higher than the *Spo11*-*1α* transcript in early zygotene anthers (Fig 2C). Unfortunately, our attempts to establish if these two isoforms have different functions were not successful.

### The *spo11-1* mutants are defective in DSB formation and homologous pairing

The SPO11 complex is required to initiate homologous recombination by creating DSBs. To analyze the presence of DSBs, we used the terminal deoxynucleotidyl transferase dUTP nick end labeling (TUNEL) assay which detects DNA damages such as ssDNA nicks and DSBs. Compared to the obvious signals detected in WT meiocytes at zygotene, most of the *spo11-1* meiocytes did not show obvious TUNEL signals. However, ∼10% of the mutant meiocytes had a few TUNEL foci (n=7/74: 9.5% for *spo11-1-Δ14,* n=2/24: 8.3% for *spo11-1-Δ03* and n=5/42: 11.9% for *spo11-1-1*) (Fig 3A, 3B). To further quantify DSB formation, we applied antibodies against γH2AX, the phosphorylated H2AX histone that is rapidly induced by DNA damage. In WT, we observed a peak in γH2AX foci during early zygotene, with an average of 518 foci per nucleus (*n*=18), whereas mutant meiocytes either lacked γH2AX signal or only showed a few discrete foci (average of 12.2 foci per nucleus, *n*=13) (Fig 3C, 3D). Similarly, some mutant cells showed very few RAD51 foci on chromatin (n=3/34: 8.8% for *spo11-1-Δ14,* n=4/37: 10.8% for *spo11-1-Δ03* and n=2/22: 9.1% for *spo11-1-1*), which is consistent with our TUNEL results (Fig S6).

**Figure 3.**
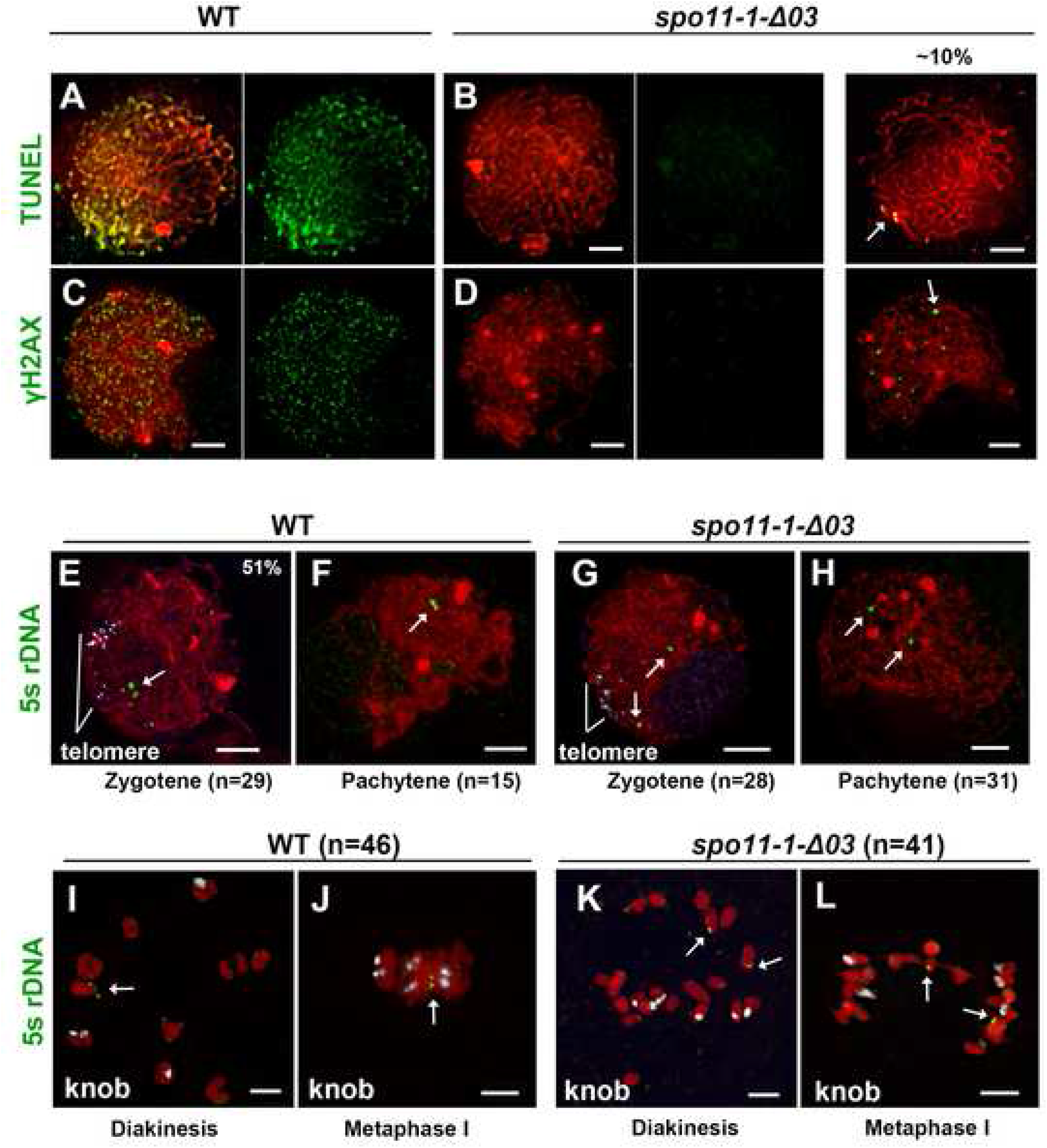
*SPO11-1* deletion affects DSB formation, chromosome pairing and homologous recombination. **(A-D)** Zygotene-stage meiocytes of WT (A, C) and the *spo11-1-Δ03* mutant (B, D) showing TUNEL (A, B), γH2AX signals (C, D) in green and DAPI stained chromatin in red. Most mutant cells exhibited no signal, but ∼10% of cells exhibited a few signals (arrows). **(E-H)** Meiocytes at zygotene and pachytene stages from WT and *spo11-1-Δ03* hybridized with 5S rDNA (green, arrows) and telomere probes (white signals in E and G) demonstrate impaired homologous pairing in the *spo11-1* mutant, although telomeres cluster normally. **(I-L)** Meiocytes at diakinesis and metaphase I from WT and *spo11-1-Δ03* hybridized with 5S rDNA (green, arrows) and 180bp-knob probes (white signals) showing that normal crossover formation is revoked in *spo11-1* meiocytes. Note that a bivalent detected in Fig 3L represents non-homologous chromosome association. All scale bars represent 5 μm.

That DNA damages, possibly including DSBs, were detected in *spo11-1* mutants, albeit at very low levels, was unexpected. Consequently, we sought to assess homologous pairing in the absence of the *Spo11-1* gene by fluorescence in situ hybridization (FISH). We found that 51% of WT meiocytes at zygotene (n=15/29) and all pachytene meiocytes (n=15) displayed paired 5S rDNA signals (Fig 3E, 3F). In contrast, among 28 zygotene-like and 31 pachytene-like *spo11-1* meiocytes, we did not detect any paired 5S rDNA signals, even though telomere clustering appeared to be normal (Fig 3G). During diakinesis and metaphase I, all WT cells (n=46) exhibited nicely paired 5S rDNA signals, whereas all *spo11-1* meiocytes (n=41) showed separate 5S rDNA signals (Fig 3I-3L). Among mutant cells, we observed a bivalent aligned at the metaphase plate with a single 5S rDNA signal, and the other 5S rDNA signal emanated from a separating chromosome (Fig 3L), suggesting that the occasional bivalents observed in *spo11-1* mutants were likely derived from non-homologous association. Taken together, these results suggest that normal DSB formation, chromosome pairing and homologous recombination is abolished in *spo11-1* mutants.

### Detection of numerous SPO11-1 foci in nuclei

To better understand SPO11-1 localization, we generated a SPO11-1 antibody against 28 amino acids at the N-terminal region and validated its specificity using peptides in the corresponding regions of SPO11-1, SPO11-2 and SPO11-3 (Fig 4C, S7). WT meiocytes from early leptotene to pachytene were subjected to immunolocalization and observed using DeltaVision deconvolution microscope (Fig 4A). At early leptotene, most SPO11-1 signals were detected around the nuclear periphery. We observed approximately 2000-3000 SPO11-1 foci in WT nuclei during late leptotene, when the chromosome mass becomes polarized. Localization of SPO11-1 in nuclei remained until the pachytene stage (Fig 4D, Table S3). In contrast, our SPO11-1 antibody detected significantly fewer foci in *spo11-1-Δ03* meiocytes (Fig 4B), similar to background levels in WT meiocytes using pre-immune serum (Fig S7C).

**Figure 4.**
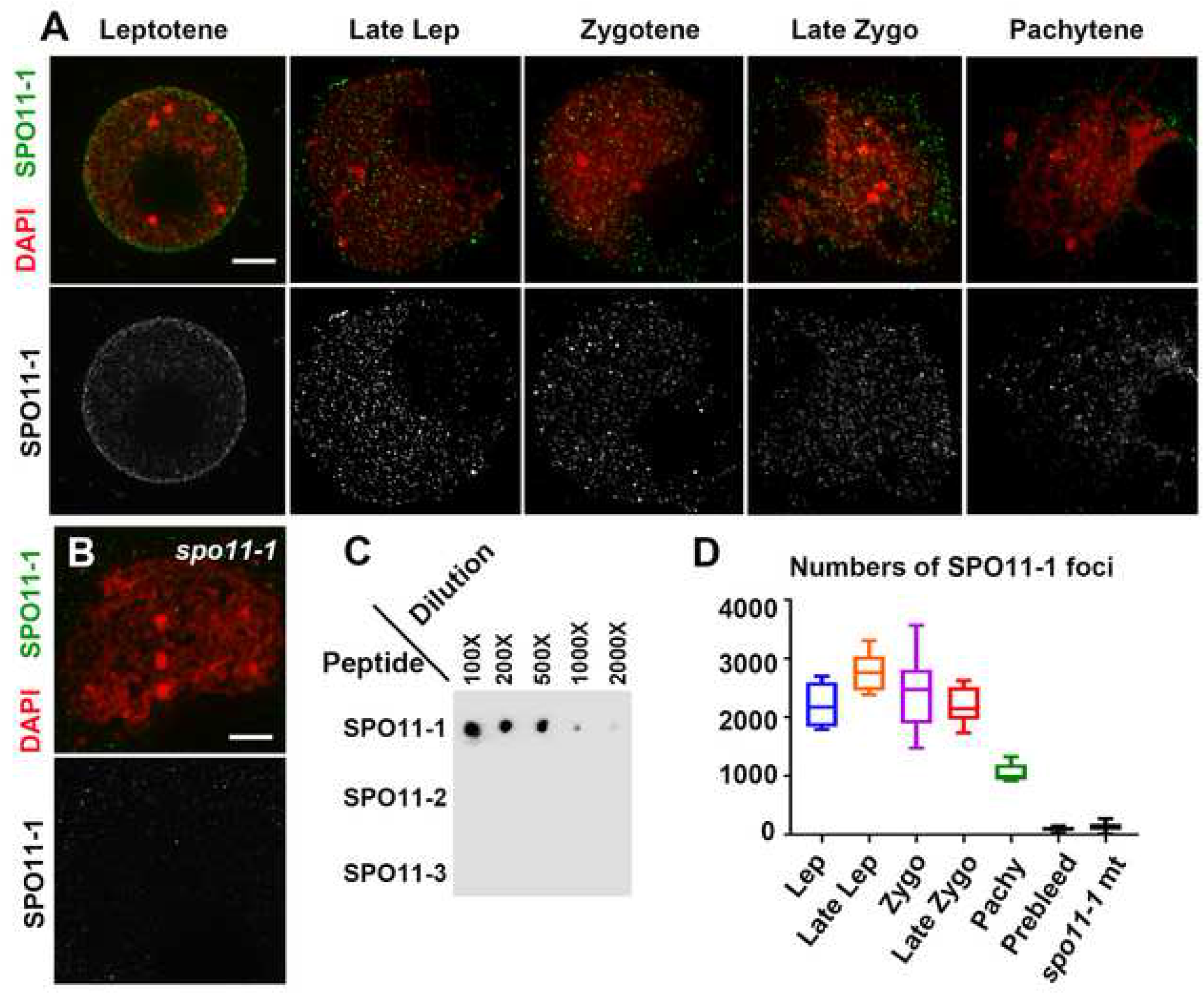
Numerous SPO11-1 signals are distributed in nuclei during early prophase I. **(A-B**) Immunostaining of SPO11-1 (green) and DAPI-stained chromosomes (red) in WT meiocytes at different stages (A) and in a *spo11-1* mutant meiocyte (B). SPO11-1 signals are shown as gray-scale in the lower panels. Scale bars represent 5 μm. **(C)** Dot blot analysis of SPO11-1 antibody against SPO11-1, SPO11-2 and SPO11-3 peptides (see Fig S7 for details). **(D)** Numbers of SPO11-1 foci detected using the affinity-purified antibody in WT nuclei at different stages and in *spo11-1-Δ03* nuclei at early zygotene. Numbers of foci detected in WT nuclei using pre-immune serum (Fig S7) are also shown (prebleed).

To further study the dynamic localizations of SPO11-1 in detail, we took advantage of the slowly progressing and well-defined meiotic stages in maize and preserved the spatial organization of meiocytes in a polyacrylamide gel for subsequent immunolocalization. Super-resolution analyses by structured illumination microscopy of snapshots of SPO11-1 distribution relative to axial protein DSY2 enabled a better depiction of DSB formation with respect to chromosome axes. In addition, the appearance of chromosomes and chromosome axes as well as the lengths of anther samples and their relative positions on a tassel provided an accurate time course for meiotic progression. In Figure 5, we show representative images of nuclei displaying SPO11-1 signals and DSY2-labeled chromosome axes from the leptotene to zygotene stages (respective montages of single Z images are shown in Fig S8-S11). In agreement with our deconvolution microscopy results (Fig 4), SPO11-1 signals were first observed mostly around the nuclear periphery and with fewer foci within nuclei (Fig 5A-5B) and were then distributed inside nuclei when the nucleolus become offset at late leptotene (Fig 5D-5E). When visualized by single Z sections (Fig 5B-C, 5E-F, 5H-I, 5K-L), SPO11-1 signals were observed as punctate foci, some of which were colocalized with DSY2-labeled AEs. To count numbers of foci, we extracted SPO11-1 signals in nuclei using a 3D Spot Segmentation algorithm and conducted quantification in the 3D ROI Manager of ImageJ [53] (Fig S12). We carefully analyzed a total of eighteen meiocytes, and again detected thousands of SPO11-1 foci distributed in nuclei (Fig 6J, Table S4). Interestingly, we recorded a wide range of SPO11-1 foci per meiocyte at each meiotic stage. In contrast, using the same detection conditions and image processing method, a significantly lower number of SPO11-1 foci was observed in *spo11-1-Δ03* mutants (Fig S13, Table S4).

**Figure 5.**
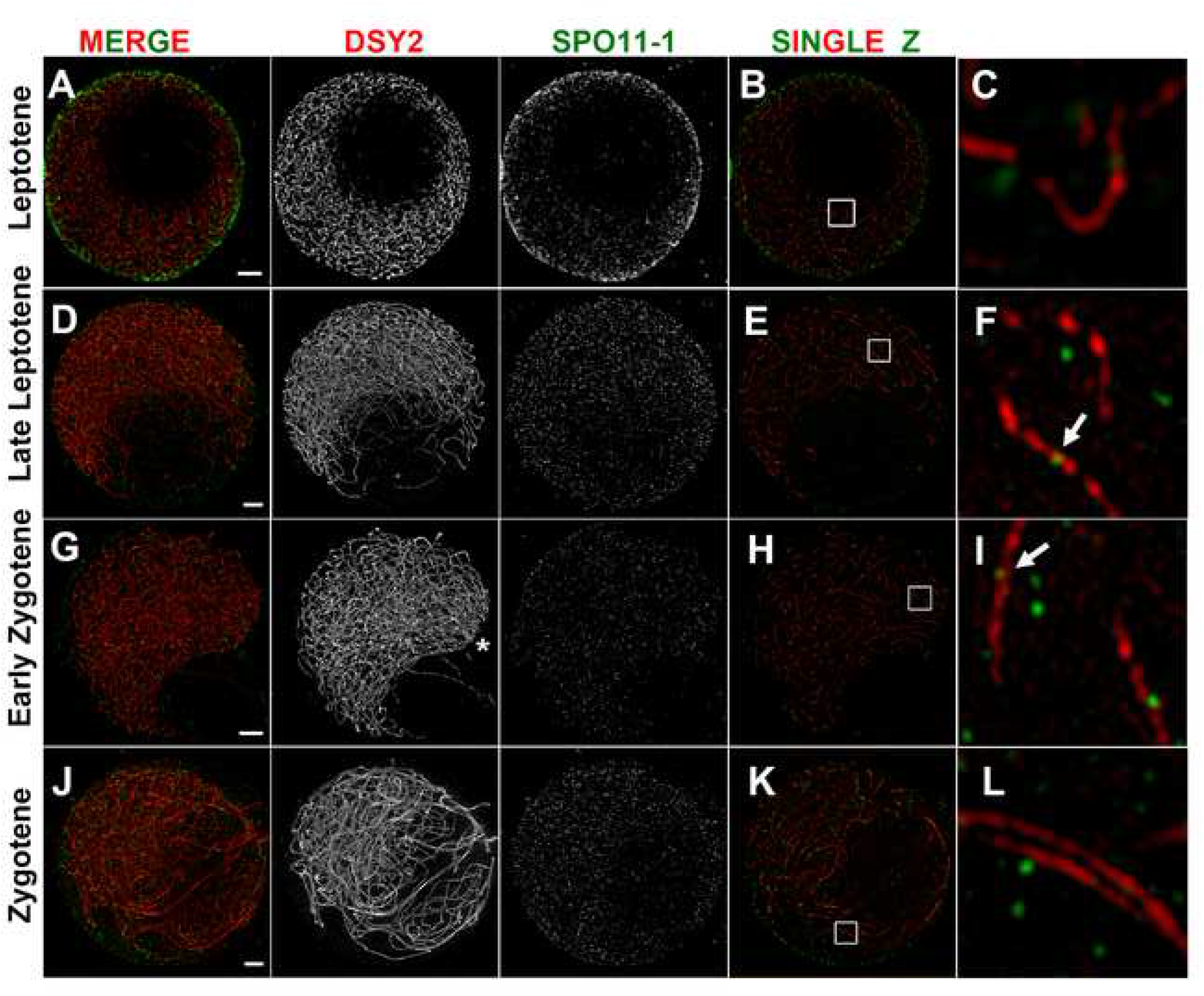
SPO11-1 localization as revealed by super-resolution microscopy in wild-type. Super-resolution images of SPO11-1 (green) and DSY2 (red) staining in WT meiocytes shown as projection images of nuclei (A, D, G, J), single Z sections (B, E, H, K), and magnified views of 2 μm^2^ regions (C, F, I, L). Scale bars represent 2 μm. Respective serial Z sections are shown in Fig S8-S11. **(A-C)** A leptotene nucleus with long and intricate chromosome axes surrounding a nucleolus exhibiting most of its SPO11-1 foci around the nuclear periphery and less signal within the nucleus. **(D-F)** A late leptotene nucleus with an off-set nucleolus exhibiting numerous SPO11-1 foci distributed within the nucleus. No obvious pre-alignment of axes was observed (Fig S9). Some SPO11-1 foci are located on DSY2-labeled AEs (arrow). **(G-I)** An early zygotene nucleus with telomere bouquet (asterisk) and pre-aligned axes (Z11-19 in Fig S10) exhibiting numerous SPO11-1 foci. Note that the nucleolus exhibits less SPO11-1 signal. **(J-L)** A zygotene nucleus with progressive pairing and synapsed regions containing numerous SPO11-1 foci.

**Figure 6.**
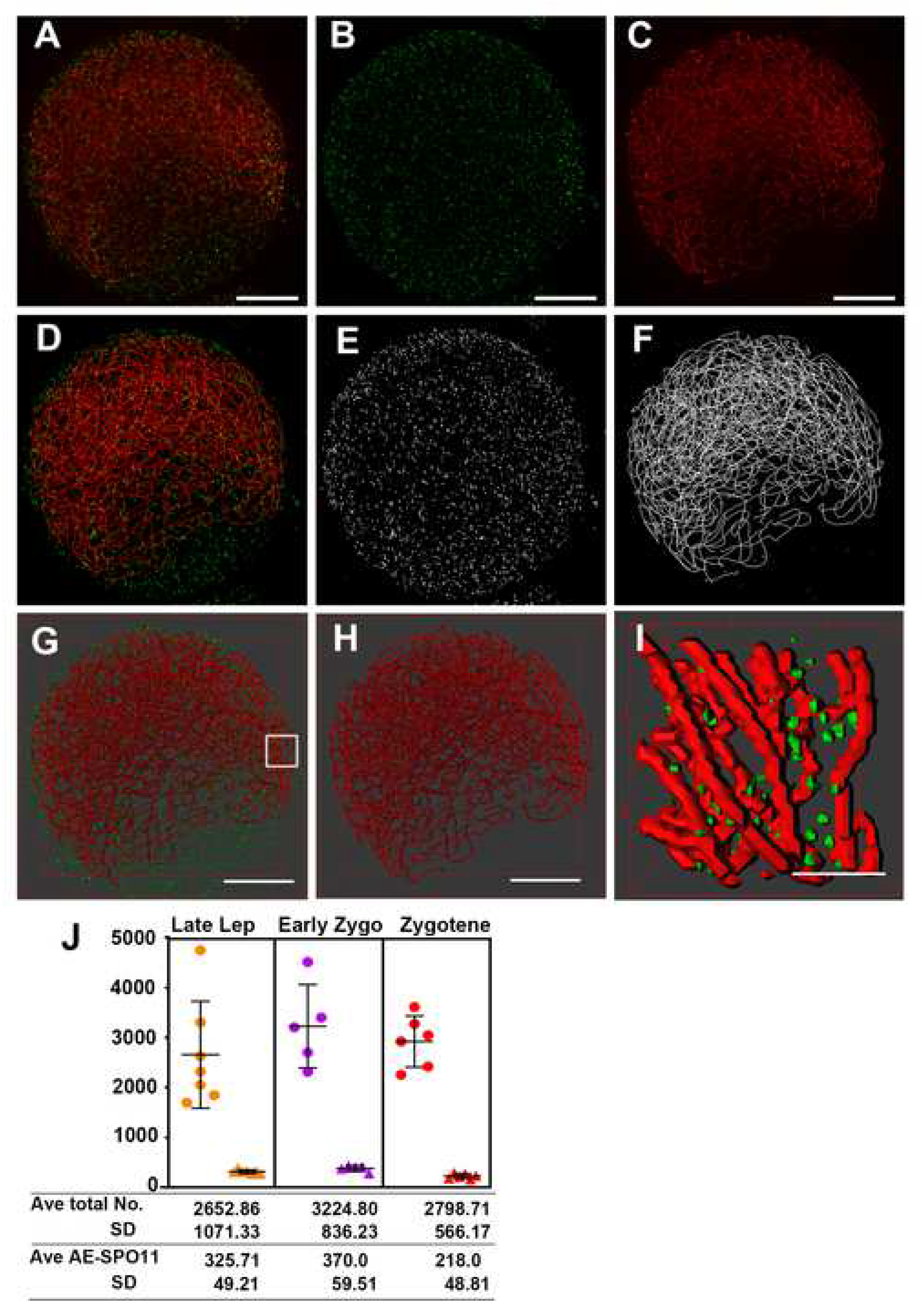
Similar numbers of SPO11-1 foci load onto axial elements. **(A-C**) Representative projection images of a WT meiocyte showing chromosome axes labeled by DSY2 (red) and SPO11-1 signals in green, captured by structured illumination microscopy. Only linear contrast adjustment was applied to enhance signals. Scale bars represent 5 μm. **(D-F)** Projection images of segmented SPO11-1 objects (E) and a skeletonized axis model (F) representing the medial lines along chromosome axes generated using our image processing pipelines (Fig S12 and S14). **(G-H)** Surface rendering images of SPO11 and DSY2, visualized by the 3D Viewer tool in ImageJ. Scale bars represent 5 μm. **(I)** A magnified 3D view of the surface rendering image in G. Scale bars represent 1 μm. A respective rotation movie is presented in Video S1. **(J)** Numbers of total SPO11-1 objects (dots) and axis-associated SPO11-1 objects (triangles) detected in meiocytes during early prophase I.

### Similar numbers of SPO11-1 foci load onto axial elements

To analyze SPO11-1 distributions relative to chromosome axes in 3D space, we developed an image processing pipeline using ImageJ [54]. The method uses the Trainable Weka Segmentation plugin, a machine learning algorithm, to segment chromosome axes (Fig S14) [55]. The Figure 6 presents DSY2 and SPO11-1 signals in a representative zygotene meiocyte that was subjected to our image processing pipelines (Fig S12, S14). After segmentation, we examined SPO11-1 objects (Fig 6E) and skeletonized DSY2-labeled chromosome axes (Fig 6F) by merging the respective images with original images and visualized them using 3D Viewer tool in ImageJ. As shown in a surface-rendered rotation movie (Fig 6I, Video S1), most SPO11-1 foci are distributed in between chromosome axes likely filled with DNA, whereas only a few SPO11-1 foci are found on chromosome axes.

To measure the physical distances between SPO11-1 foci and the nearest chromosome axis, we computed the skeletonized axis model representing medial lines along chromosome axes in 3D stacks and generated a 3D distance map which assigns propagating distances from each point along axes to neighboring voxels in the 3D space (Fig S15). By applying segmented SPO11-1 objects onto the corresponding 3D distance map, we obtained the geodetic distance between each SPO11-1 object and the closest DSY2-labeled axis (Fig S16, Table S5). In Figure S17, we display the distribution of these minimal distances in each meiocyte and show that a majority of SPO11-1 objects was located within a radius of 0.6 μm from a neighboring axis. More than 50% of the foci in each meiocyte were distributed within DAPI-stained chromatin regions (a radius of 0.3-0.4 μm) in most of the cells examined. We defined SPO11-1 signals as AE-associated objects when the minimal distance to the closest chromosome axis was zero (Table S5). Surprisingly, despite a wide range (∼1691-4750) of SPO11-1 foci being distributed in nuclei, approximately 300-400 SPO11-1 foci were associated with chromosome axes during recombination initiation and those numbers decreased slightly during zygotene (Fig 6J, Table S4).

### Maize *spo11-1* mutants present an aberrant axial element structure and promiscuous synapsis

To better characterize phenotypes of chromosome axes in the absence of *Spo11-1*, we compared AEs of WT and mutant meiocytes sampled from similarly sized anthers by DSY2 immunostaining. Our results show that WT and mutant meiocytes at leptotene possessed long and curly AEs enclosing a large, centered nucleolus (Fig 7A, 7B). At late leptotene, the chromosome reorganization that offsets the nucleolus position occurs normally in *spo11-1* mutants (Fig 7C, 7D). In WT, pairing and synapsis take place mostly from telomere clustering regions and extend toward interstitial regions, often manifesting as multiple aligned bands of bright DSY2 signal intensity (Fig 7E). However, in contrast to WT meiocytes in which AEs became smoother and more linear, we observed curly and aberrant AEs with uneven DSY2 accumulation in *spo11-1* mutants (compare Fig 7C, 7E, 7G and Fig 7D, 7F, 7H). The erratic structure of these AEs become more evident when we visualized the projection images of a few Z-sections (see magnified views in Fig 7A-H, Fig S18). To describe and quantify the cytological aberrance of these AEs, we measured the total lengths of chromosome axes (Fig S14) and also computed their curvatures [56] (Fig S19). In Figure 7I, we show that AE length diminished as meiosis progressed in both WT and mutant meiocytes (Table S6). Notably, AEs in WT meiocytes shortened from ∼4 to ∼3 mm during the leptotene to late leptotene stages before synapsis took place, but a corresponding curtailment was not found in mutant meiocytes. Moreover, kappa values of AE curvature in mutant cells were elevated in late leptotene and zygotene stages relative to WT (Fig 7J), suggesting that not only is compactness of AEs affected, but their morphology is also altered, appeared as more flexuous axes in *spo11-1* mutants.

**Figure 7.**
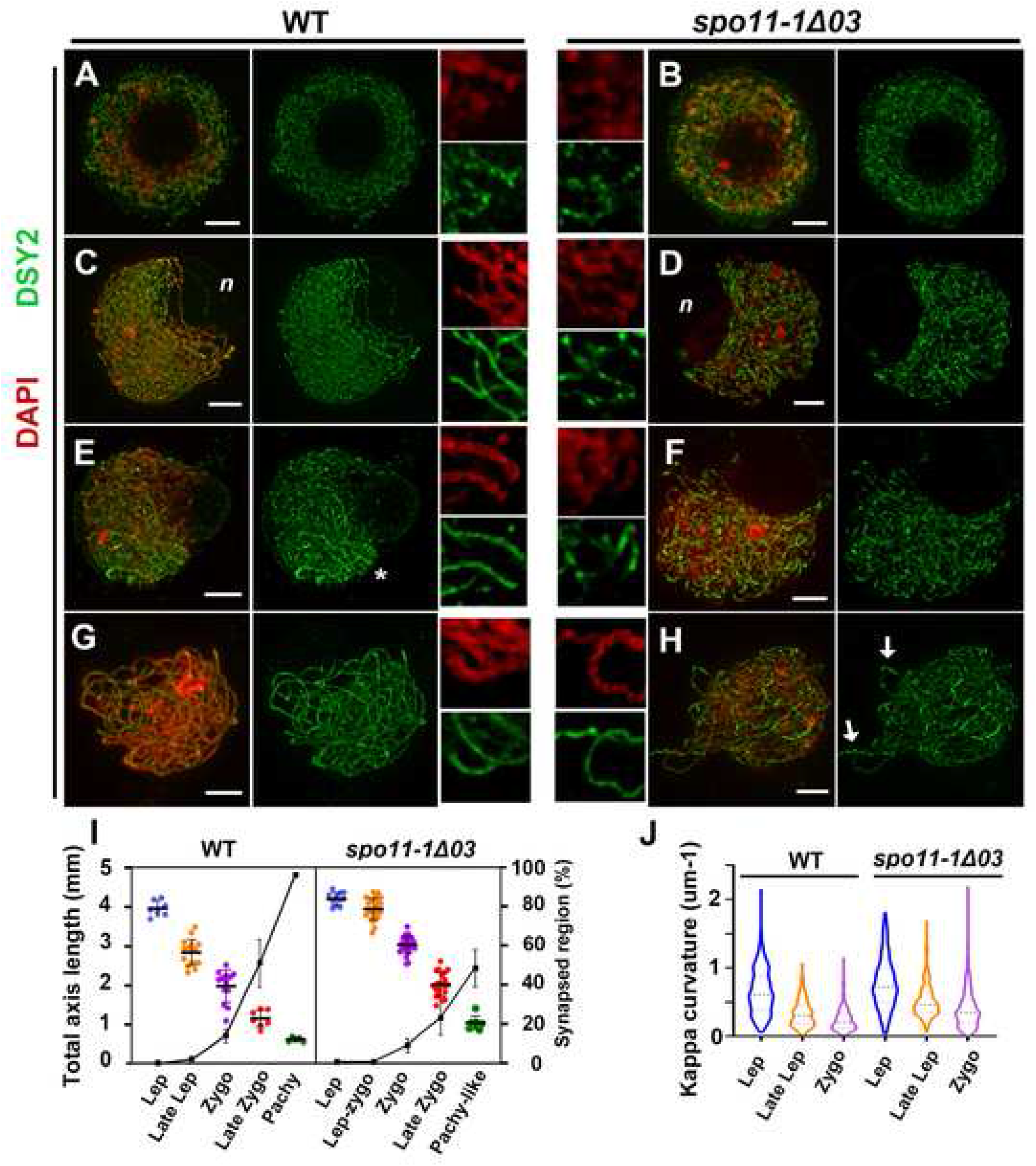
An aberrant axial element structure in *spo11-1* mutants. Immunostaining of DSY2 (green) and DAPI staining (red) in WT (A, C, E, G) and *spo11-1-Δ03* meiocytes (B, D, F, H). Maximal projection images are displayed for entire nuclei. Magnified images of 5 μm^2^ regions are shown in the middle panel. Scale bar represents 5 μm. **(A, B)** The morphology of chromosome axes is similar in WT and mutant meiocytes when DSY2 first appears in the nucleus at the early leptotene stage. **(C, D)** At late leptotene, AEs appear progressively as linear forms in WT (C). In contrast, abnormal AE structures are observed in mutant meiocytes (D). Note that DSY2 signals intensity is uneven in mutant meiocytes, with atypically bright fluorescent patches, unlike the uniform signals in the WT. The off-set nucleolus is marked with *n*. **(E-F)** Synapsis, manifested as paired doubled DSY2 stretches, is observed primarily near the telomere bouquet (*) of the WT (E), which is absent from mutant meiocytes (F). **(G-H)** A late zygotene WT meiocyte (G) showing consistent DSY2 signal along chromosomes, unlike the unevenly accumulated DSY2 (arrows) in *spo11-1* meiocytes (H). **(I)** Dot plot graph showing total lengths (mm) of DSY2 signals from the leptotene to pachytene stages in WT and *spo11-1* meiocytes. Each dot represents a measurement from one meiocyte. The linear plot represents the percentage of synapsed regions, as measured by ZYP1 signal length over total DSY2 signal length. **(J)** Distribution of kappa values representing AE curvature, as measured from five meiocytes for each stage in WT and *spo11-1* mutants.

Super-resolution images of ASY1 and DSY2 staining in WT meiocytes revealed diminished ASY1 intensity and enhanced DSY2 intensity in synapsed regions, which reflects the progression of synapsis (Fig 8A). In mutant meiocytes, DSY2 and ASY1 both exhibited a similar abnormality (Fig 8B), seemingly due to incorrect patterning of axial proteins. Similarly, curly and unevenly accumulated ASY1 staining were observed in *spo11-1* mutants (Fig 8C, 8D). Despite the aberrant AEs and the lack of DSBs, all maize *spo11-1* mutants showed delayed synapsis as short stretches of ZYP1 signal appear later than those observed in WT (Fig 9C, S20). In contrast to WT at pachytene, mutants displayed discontinuous ZYP1 bands in tangled chromosomes, likely between non-homologous chromosomes (Fig 9D). Transmission electron microscopy of silver nitrate stained SCs revealed some synapsed regions with fold-back conformations, suggesting promiscuous synapsis (Fig 9G, S21). Together with our FISH results (Fig 3H, 3L), we conclude that synapsis is initiated randomly between non-homologous chromosomes in the *spo11-1* mutants.

**Figure 8.**
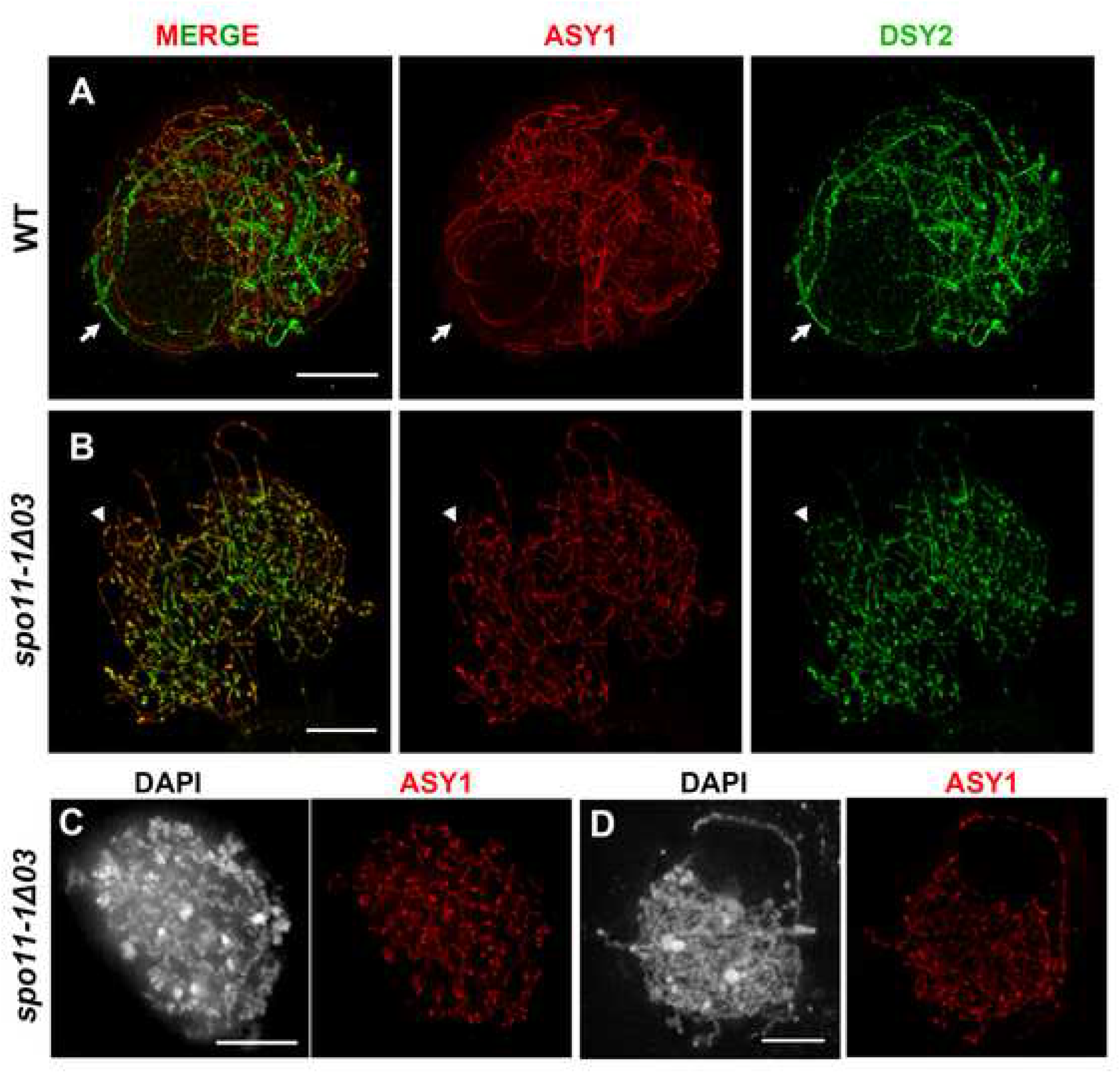
Axial element proteins ASY1 and DSY2 exhibit abnormal localization in *spo11-1* mutants. **(A, B)** Zygotene meiocytes of the WT (A) and *spo11-1* mutant (B) immunolabeled with ASY1 (red) and DSY2 (green), and observed by super-resolution microscopy. Note that synapsed regions in the WT (arrows) exhibit diminished ASY1 and enhanced DSY2 signals. In mutant meiocytes, both ASY1 and DSY2 accumulate abnormally (arrowheads). **(C, D)** Abnormal chromosome axes with curly and uneven ASY1 staining are observed in *spo11-1* meiocytes. Scale bars represent 5 μm.

**Figure 9.**
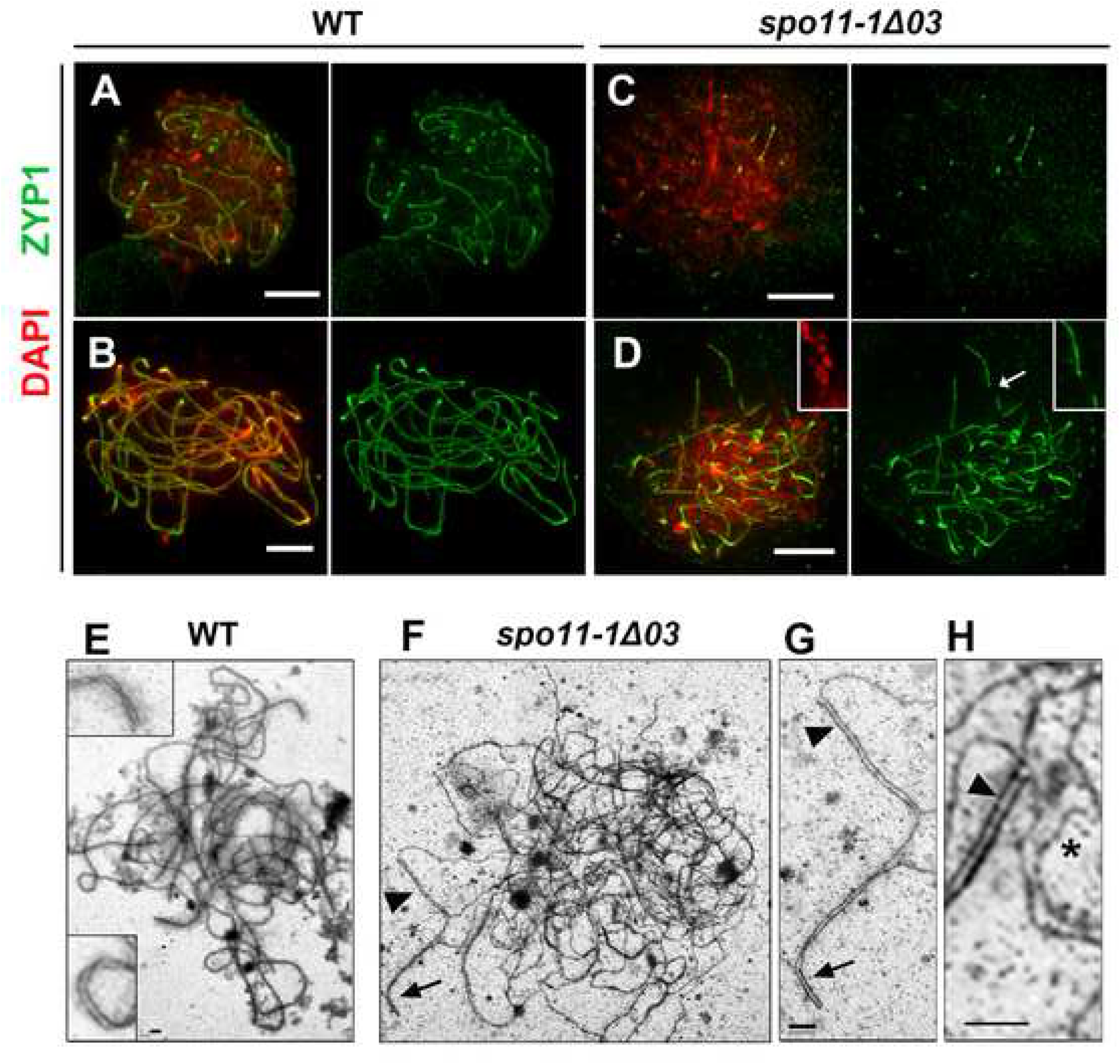
Transverse filament loading is initiated but is not completed. **(A-D)** Immunostaining of ZYP1 (green) with DAPI staining (red) in WT (A, B) and *spo11-1-Δ03* (C, D) meiocytes at zygotene (A, C) and pachytene stages (B, D). Note the magnified region from the pachytene-like mutant meiocyte (D) showing discontinuous ZYP1 stretches (arrow). Scale bars represent 5 μm. **(E-H)** Transmission electron microscopy of synaptonemal complex spreads of WT (E) and *spo11-1-Δ03* (F-H) meiocytes. Unlike the complete synapsis in WT, mutants exhibit abnormal synapsis (G and H). Note that synapsis occurs within one axis, forming “fold-back” structures (arrow), and promiscuous synapsis can also be seen as unequal synapsis with an overhang (arrowheads). In addition, unsynapsed AEs in mutants appear as curly and twisted structures (asterisk in H). Scale bars represent 1 μm.

## Discussion

Molecular and genetic studies over past decades have garnered substantial information on meiotic DSB formation. First, programmed DSBs in meiosis require SPO11 and the meiotic topoisomerase VI subunit B protein, forming a complex [57], as well as other DSB-accessory proteins [40]. Second, DSBs form preferentially in open chromatin regions residing within chromatin loops that emanate from chromosome axes [58]. Third, potential meiotic DSB sites are brought to chromosome axes [40, 59, 60]. Fourth, the SPO11-containing complex is activated and then generates a DSB during or after the loop-tethering event [1]. These findings favor the ‘tethered loop-axis complex’ model and also explain the observation of a hierarchical combination of factors governing DSB sites. Although much is known about this process, a full picture of is still elusive.

One aspect that remains unclear is how DSBs form in relation to SPO11 localization and chromosome axes. It is difficult to detect SPO11 in most organisms, probably due to its narrow timeframe of expression, its dynamic localization, and/or low sensitivity for *in situ* detection of its protein molecules. It has been observed by immunocytochemistry of chromosome spread samples in *S. cerevisiae* [61], *S. macrospora* [62], *Arabidopsis* [63] and mouse [64]. These results show that SPO11 foci appear on chromosome axes and/or in chromatin regions during the leptotene to early pachytene stages. Unfortunately, due to limitation of microscope resolution and slide preparation of chromosome spreads, it seems difficult to analyze SPO11 localization in detail.

However, due to the remarkable advantages of maize for cytological observations, our analyses have led to several interesting observations that provide new insights into DSB formation. By super-resolution microscopy and sophisticated image processing on 3D-preserved maize meiocytes, we have uncovered dynamic SPO11-1 localization during recombination initiation. First, SPO11-1 signals were observed mostly around the nuclear periphery and later distributed in nuclei. Next, although a broad range of thousands of SPO11-1 foci was detected in nuclei with more than half of them colocalized with DAPI-stained chromatin, only relatively similar numbers (∼350) of SPO11-1 foci were associated with chromosome axes (Fig 6). It is intriguing that we detected a large amount of SPO11 distributed around or within chromatin regions since the SPO11 enzyme is intrinsically a threat for genome integrity. However, this finding may reflect the fact that SPO11 localization on chromatin loops alone is inadequate to make DSBs. Moreover, previous studies have suggested that 200-500 DSBs are generated during each round of maize meiosis [65], which is in accord with numbers of AE-associated SPO11-1 signals detected in this study, implying that only AE-associated SPO11 foci can generate DSBs. In addition, the timing of SPO11-1 localization on chromosome axes coincides with the leptotene to zygotene transition when homologous alignment and telomere clustering are taking place. Taken together, our results suggest a model whereby numerous SPO11 proteins bind on potential DSB sites, and the competitive and dynamic tethering events that link SPO11-marked hotspots to chromosome axes may play a critical role in DSB formation. It is not clear if the SPO11-1 foci on chromatin represent DSB hotspots in our analyses. However, in support of this notion, a previous study in *S. cerevisiae* using a standard ChIP protocol has detected a transient and noncovalent association of SPO11 with meiotic hotspots [61]. That transient association is independent of DSB formation since it was also detected in *spo11-Y135F* mutation, suggesting that SPO11 accumulates transiently at hotspots before stable DSBs form at some of sites. It is also interesting to note that maize SPO11-1 signals remained until the pachytene stage which is consistent with previous finding of detecting SPO11-1-myc foci in the Arabidopsis SPO11-1-myc *spo11-1* mutant line [63].

In our study, we have characterized the phenotype of maize *spo11-1* mutants in detail and showed that loss of *Spo11-1* in maize results in meiotic sterility, as has been observed in other species. Unexpectedly, we also observed abnormal chromosomal axes in maize *spo11-1* mutants. First, leptotene AEs in mutants were indistinguishable from those of WT in terms of their length, morphology and curvature. However, differences became apparent when meiosis proceeded to late leptotene when a global re-organization of chromosome configuration takes place. Our results show that *spo11-1* meiocytes displayed longer chromosome axes with aberrant and uneven accumulation of DSY2 and ASY1 patches, unlike the smooth and shortened chromosome axes of WT meiocytes. Moreover, at early zygotene when synapsis is initiated shortly after pre-alignment in WT, mutant cells showed delayed SC nucleation and aberrant chromosome axes. The curly AEs in *spo11-1* meiocytes may result from mis-patterning of axial element proteins, which appear after the late leptotene stage. This observation points to the existence of a possible mechanism that is established in response to DSBs as a part of AE maturation process. For example, when DSBs are made in recombinosomes that are located on chromosome axes, a localized structural transition may be triggered in the chromosome axes such as by remodeling axial proteins [66], by phosphorylation of ASY1/HOP1 and REC114/PHS1 via the ATM-mediated DNA damage response [15, 67], or simply by AE condensation, to prepare the sites for sequential interhomolog recombination or to prevent additional DSB generation by rejecting further tethering events. In the absence of SPO11-1 (or due to failure of DSB formation), these responses are not induced, so the chromatin and chromosome axes probably remain in a DSB susceptible state. Nevertheless, other meiotic cell cycle modules, such as cyclin-dependent kinase, may still stimulate meiotic progression to ultimately relax the barrier of SC assembly, resulting in synapsis between non-homologous chromosomes in pachytene-like mutant meiocytes.

It has been reported that maize meiocytes during the leptotene-zygotene transition (prezygotene) undergoes dramatic changes, including increased total chromosome volume, decondensed heterochromatic knobs, offset-localized nucleolus and clustering of telomeres [41]. Meiotic DSBs are thought to form at this stage or shortly prior to this transition [68]. Here, we have shown that during this transition the numbers of highly copious SPO11-1 foci inside the nucleus are variable, whereas relatively similar numbers of SPO11-1 foci are loaded on AEs. Interestingly, AEs in wild-type cells gradually became smoother and more linear, which does not occur in *spo11-1* meiocytes at the same stage. In addition, the shortening of AEs (from 4 to 3 mm) that we detected in wild-type meiocytes at this transition is likely a response to the presence of SPO11 or DSB formation, but it is not caused by synapsis since ZYP1 signals were detected at later stages in WT meiocytes.

To date, programmed DSB formation in eukaryotic meiosis has been shown to require SPO11. Unlike the single SPO11 protein in mammals and fungi, plants encode at least three SPO11 proteins [69]. In Arabidopsis, genetic analyses revealed that two non-redundant SPO11 proteins are required for meiotic recombination [70–72], suggesting that DSBs are catalyzed by a SPO11-1/SPO11-2 heterodimer. Maize has three SPO11 related proteins. Although the meiotic functions of maize SPO11-2 and SPO11-3 are not clear, loss of SPO11-1 alone severely affects DSB formation and homologous recombination. Despite a few DBSs being detected in *spo11-1* mutants by our TUNEL assay and γH2AX, RAD51 immunolocalization, these DSBs (or other types of DNA damages) was not repaired by homologous recombination, suggesting that neither SPO11-2 nor SPO11-3 possesses redundant functions for SPO11-1.

Current DSB maps generated by Spo11-oligo sequencing have provided a landscape of recombination initiation sites [63, 73–75]. However, despite thousands of DSB hotspots having been identified, only a few hundred DSBs are usually detected during each round of meiosis. Considering the probabilistic nature of DSB formation, further investigations of DSB regulation, distribution and timing at the level of individual cells will likely fill this information gap. Here, our observations based on super resolution microscopy have revealed for the first time in any organism the dynamic localization of SPO11-1 relative to AEs, providing evidence for the ‘loop-tethered axis’ model. By characterizing the phenotypes of maize *spo11-1* mutants, we suggest that there is an intimate relationship between DSB formation and the structure of chromosome axes.

## Material and methods

### Maize material

Maize inbred lines B73, A344, A632 and Mo17, as well as wild type siblings of the *Mu* insertional mutants and *spo11-1* mutant alleles, were grown in the greenhouse of Academia Sinica and previously in the UC Berkeley greenhouse and experimental field. The *mtm99-14 / spo11-1-Δ14* and *mtm00-03 / spo11-1-Δ03* mutants came from two independent genetic screens for maize meiotic mutants using a Mutator (*Mu*) population, respectively, during the 1999 and 2000 field seasons [47, 48]. The *spo11-1-1* line was found in the TUSC reverse genetic resource [52].

### Genetic mapping and *Mu* insertion determination

To map the *mtm00-03* mutation, bulk segregation analysis (BSA) of two *mtm00-03* F2 populations created by crossing the *mtm00-03* line with inbred A344 or B73 were used, as previously described [76]. Sterile plants showed tight linkage to the polymorphic markers IDP8472 and particularly IDP4279 that map to the end of the short arm of chromosome 5 on Bin 5.01 between physical positions 5.3 and 8.5 Mb. For each mutant, *Mu* insertion border sequences that are associated with segregation of phenotype were determined using the *Mu*-Illumina border determination approach, as previously described [51].

### RT-PCR analysis

RT-PCR analyses were conducted on RNA extracted using Trizol from meiotic anthers, as described previously [77]. The *Afd1* 252 base pair (bp) cDNA and *PhyC* 444 bp cDNA fragments, as well as the 3’ and 5’ portions of *Spo11-1* transcripts, were amplified using specific primers (Table S7). The *PhyC1* and *PhyC2* fragments presented as two distinct bands from genomic DNA but with an identical size to cDNA. All PCRs were performed using DMSO and the Invitrogen GoTaq kit with 30 cycles using annealing temperature of 52 **^°^**C except for the SPO11-1 5’ amplification using 60 **^°^**C. Quantitative PCR of *Spo11-1α* and *Spo11-1β* transcripts was performed using FastStart Universal SYBR Green Master mix with specific primers (Table S7). Cyanase was used as an internal normalization control. PCRs were performed on an ABI 7500 real-time PCR system in a 64-well plate. The RT-qPCR reaction was performed in triplicate for each RNA sample. Specificity of the amplification products and Ct values were analyzed using ABI prism dissociation curve analysis software.

### Antibody production

To generate anti-SPO11-1 antibody, we used a peptide corresponding to amino acids 9-36 of SPO11-1 protein (RAAPLEGDEQQLRRRLEEAALLLRRIKG) as antigen to generate a rabbit polyclonal antibody, followed by affinity purification (LTK BioLaboratories, Taiwan). Antibody specificity was validated by dot-blot analysis (Fig S7).

### Western blot

Anthers were dissected from immature tassels after their meiotic stages and sizes had been determined and were then snap frozen in liquid nitrogen. Protein extraction was performed as previously described [37]. Two hundred micrograms of proteins were loaded in a 20 cm long, 10% SDS-PAGE gel, run for 18 h at 100 V, and then transferred to a polyvinylidene fluoride (PVDF) membrane. The PVDF membrane was blocked overnight with 5% milk in Tris-buffered saline-Tween (TBST) solution (containing 1X TBS and 0.1% Tween 20) at 4 °C. Subsequently, the PVDF membrane was incubated with anti-SPO11-1 antibody (1:1000 dilution in 5% milk/TBST) and then with horseradish peroxidase conjugated goat anti-rabbit IgG secondary antibody (1:5000 dilution in TBST). After a TBST wash, signals were detected using UVP Biospectrum 600 imaging system.

### Immunolocalization

Immature tassels were carefully removed from 4- to 6-week-old plants and kept in moist paper towel until dissection or fixation. Anthers were dissected after their meiotic stages and sizes had been determined and their positions on a tassel were recorded as a relative time course. We analyzed meiosis in synchronized cohorts of meiocytes in anthers using characteristics described previously [78]. Immunostaining was performed as described [79] with primary antibodies anti-DSY2 (1:200) [37], anti-ASY1 (1:400) [33], anti-γH2AX (1:200) [80], anti-RAD51 (1:100) [81], anti-AFD1 (1:200) [42], anti-ZYP1 (1:100), and anti-SPO11-1 (1:100). The anti-RAD51 antibody was a gift from Wojciech Pawlowski (Cornell University) and anti-γH2AX was a gift from Hong Ma and Yingxiang Wang (Fudan University). Conjugated secondary antibodies were obtained from Molecular Probes. Slides were imaged using either an LSM microscope 780 with ELYRA equipped with SR-SIM (Zeiss) or a Deltavision core microscope (GE).

### Image processing

The super-resolution z-stacking images were examined using ZEN2012 software (Zeiss). All image processing procedures were carried out in “ImageJ-FIJI” software. The SPO11-1 signals were segmented as described in Fig S12. Chromosome axes were skeletonized as described in Fig S14. To measure distances between SPO11-1 and chromosome axes, a 3D distance map of DSY2-labeled chromosome axes was created using the plugin “3S Image Suite” (Fig S15), and 3D measurement of objects and distance information was analyzed using the “3D ROI Manager” tool in 3D Image Suite (Fig S16). The curvature of chromosome axes was analyzed using the Kappa plugin (Fig S19). All super-resolution images used for measurement have been deposited in the BioStudies database (http://www.ebi.ac.uk/biostudies) under accession number S-BSST322.

### TUNEL assay and TEM

The TUNEL assay for DSB detection was performed as described previously [82]. First, meiocytes were embedded in polyacrylamide pads as described in the immunostaining protocol above. The TUNEL reaction was performed in a humid chamber for 2.5 h at 37 °C with the *in Situ* Cell Death Detection Kit (Roche) using 100 μL of reaction mixture per slide. Slides were mounted and imaged with a Deltavision core microscope. TEM analysis was performed as described previously [82, 83].

## Author Contributions

JCK and AR designed and carried out field experiments, molecular assays and immuno-cytological analyses. IG did the original forward genetic screen, determined the allelism between *mtm00-03* and *mtm99-14,* and conducted initial cytological analyses of the *spo11-1* mutants. DHL and CTW performed super-resolution microscopy and analyzed data. LT did the initial BSA mapping, made crosses, determined the allelism between *mtm00-03* and *mtm99-14,* conducted the TEM analysis, and provided the DNA of the *mtm00-03* mutants for *Mu*-Illumina sequencing. AKGA did the RT-PCR analysis. YHK performed quantitative PCR for alternative spicing. KK did the BSA mapping and the RT-PCR analysis. RWC and AB did the *Mu*-Illumina border sequencing for *mtm99-14* and *mtm00-03*. RM provided the original seed stocks of the TUSC lines (*spo11-1-1*). ZC initiated the project. CJRW wrote the paper and supervised the project.

## Database Information

All super resolution microscopic images are available in the BioStudies repository (http://www.ebi.ac.uk/biostudies) under accession number S-BSST322.

## Acknowledgments

We thank Jihyun Moon for advice on BSA mapping analysis. We thank Sidae Lee and Angel Jung for help with specimen collection for DNA sampling, and Patricia Elda Rueda for plant genotyping. We appreciate Wojtek Pawlowski, Hong Ma and Yingxiang Wang for their RAD51 and γH2AX antibodies. We thank Shu-Chen Shen (Scientific Instrument Center, Academia Sinica) for help with super-resolution imaging. Special thanks to Thomas Boudier (Walter& Eliza Hall Institute, Australia; now a visiting scholar in Academia Sinica) for his tremendous helps with image analysis. Mathilde Grelon provided valuable suggestions on this manuscript. This research was supported by a grant from NIH GM48547 (to WZC), a research grant from UNAM PAPIIT IA201217 (to AR), a grant IOS-1339130 from the US National Science Foundation (to AB), and a grant (to CJRW) from the Ministry of Science and Technology, Taiwan (107-2923-B-002-001-MY4).

## Supporting Data

**Video S1.** A rotation movie of the surface-rendered image shown in Figure 6I that has been magnified from a labeled region in Figure 6G showing the relative 3D positions of chromosome axes (red) and SPO11-1 signals (green).

**Table S1. Genetic segregation of male sterility for *spo11-1* mutant alleles and progeny of the allelism test.**

**Table S2. Female fertility shown by seed sets of cobs pollinated with WT Mo17 pollen.**

**Table S3. Numbers of SPO11-1 foci detected in WT and *spo11-1-Δ03* meiocytes.**

**Table S4. Numbers of SPO11-1 foci detected by structured illumination microscopy.**

**Table S5. Geodetic distance of SPO11-1 foci to the nearest DSY2-labeled axes.**

**Table S6. Total length of chromosome axes during early meiosis in WT and spo11-1 mutant meiocytes.**

**Table S7. Primers used in this study.**

---

**Figure S1.**
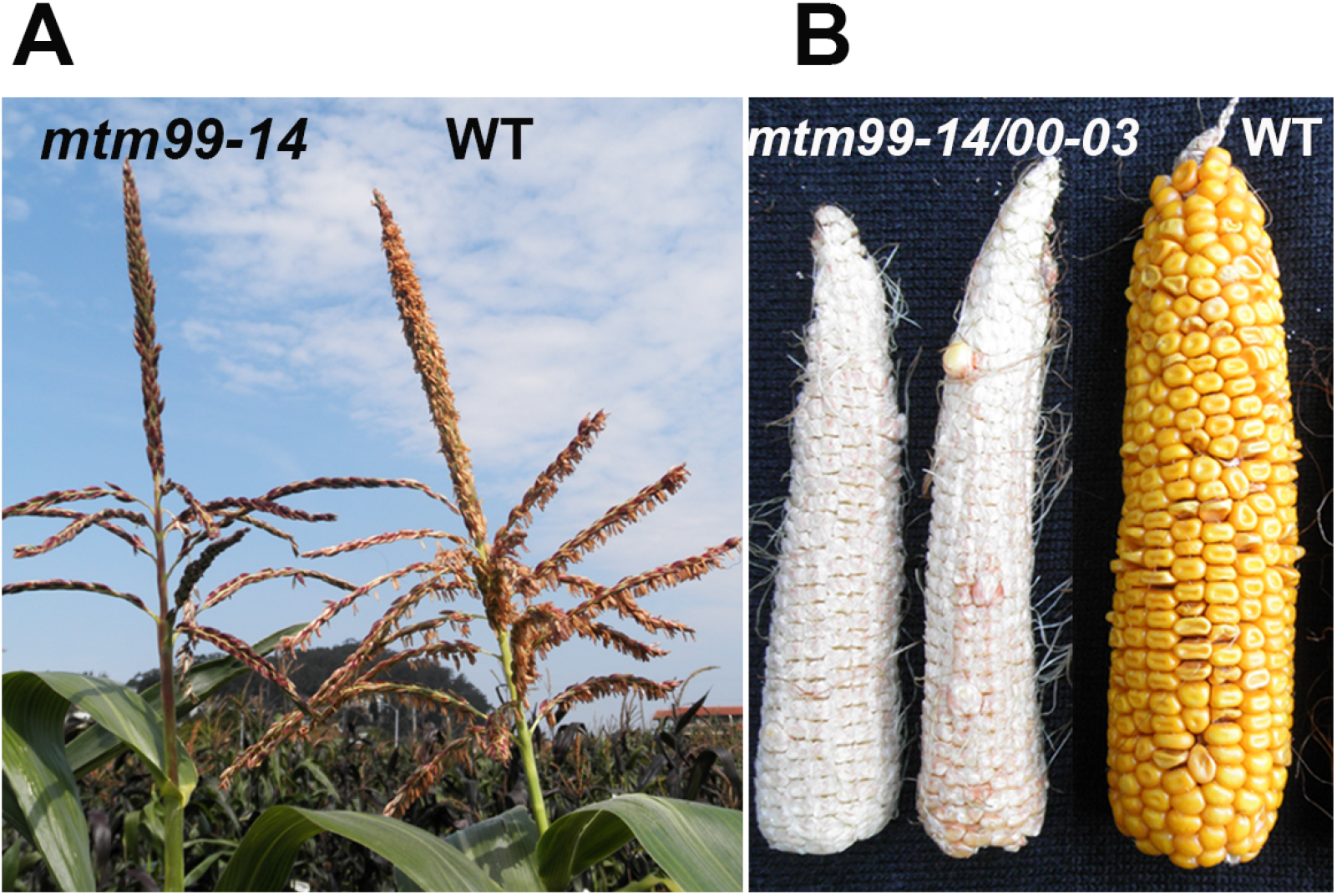
Representative examples of male and female phenotypes observed in mutant maize plants. **(A)** Male inflorescence (tassel) of the *mtm99-14* mutant and wild type (WT) at time of pollen shedding. The WT plant exhibits protruding anthers and shed pollen. Protruding anthers are very rare in mutant plants and they do not shed pollen grains. **(B)** The two maize ears on the left are from heteroallelic *mtm99-14/mtm00-03* plants pollinated with WT (Mo17) pollen and mainly present shriveled ovules with few kernels. The ear on the right is from a WT sibling plant that is full of kernels.

**Figure S2.**
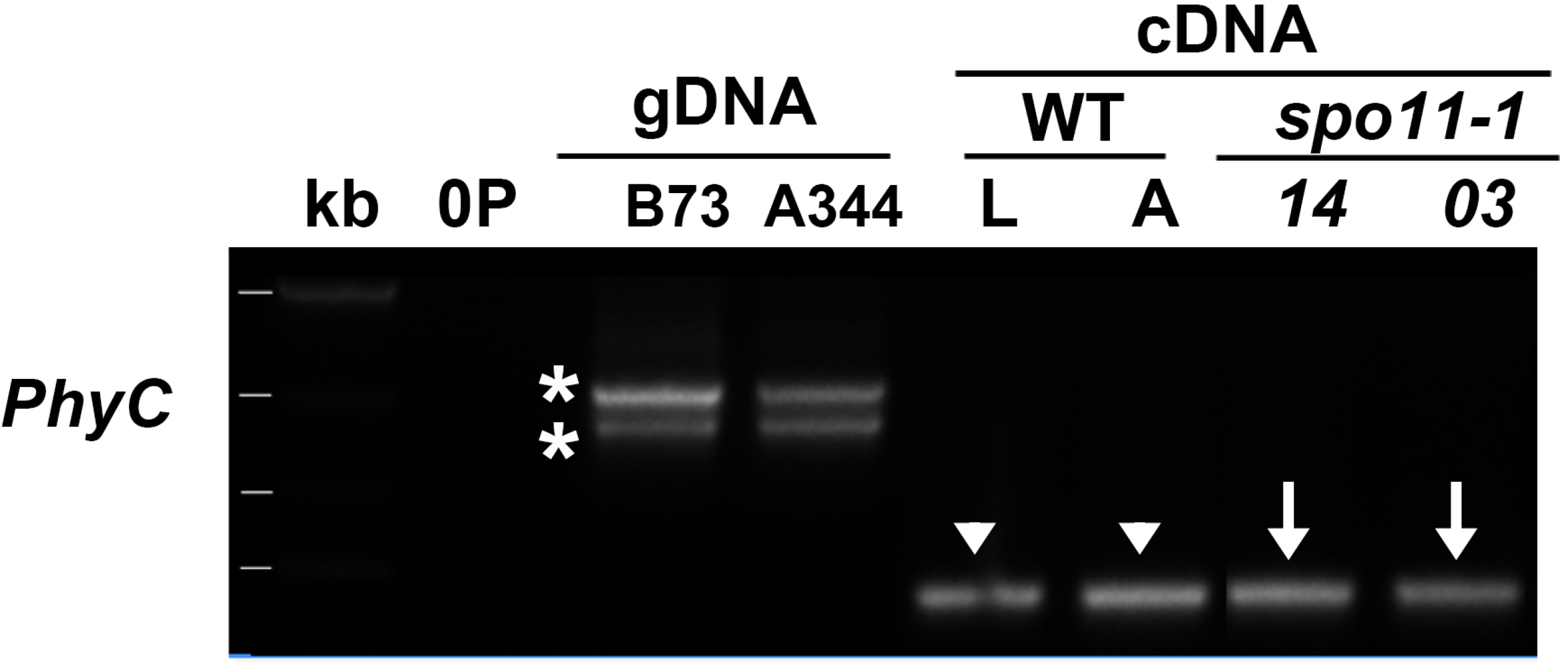
Amplification of *PhyC* genes from WT and *spo11-1* mutant alleles. Given 94% identity between the *PhyC1* and *PhyC2* coding regions, both *PhyC1* and *PhyC2* fragments (asterisks) were amplified from genomic DNA (gDNA) of B73 and A344 WT using primers AR105/AR106 (Table S7). 0P represents a negative PCR control lacking DNA template. By RT-PCR, the same primers amplified both transcripts (arrowheads) of identical size from WT leaf (L) and anther (A). However, sequencing results confirmed that only the *PhyC1* transcript (arrows) was detected in the *mtm99-14* (*14*) and *mtm00-03* (*03*) mutants.

**Figure S3.**
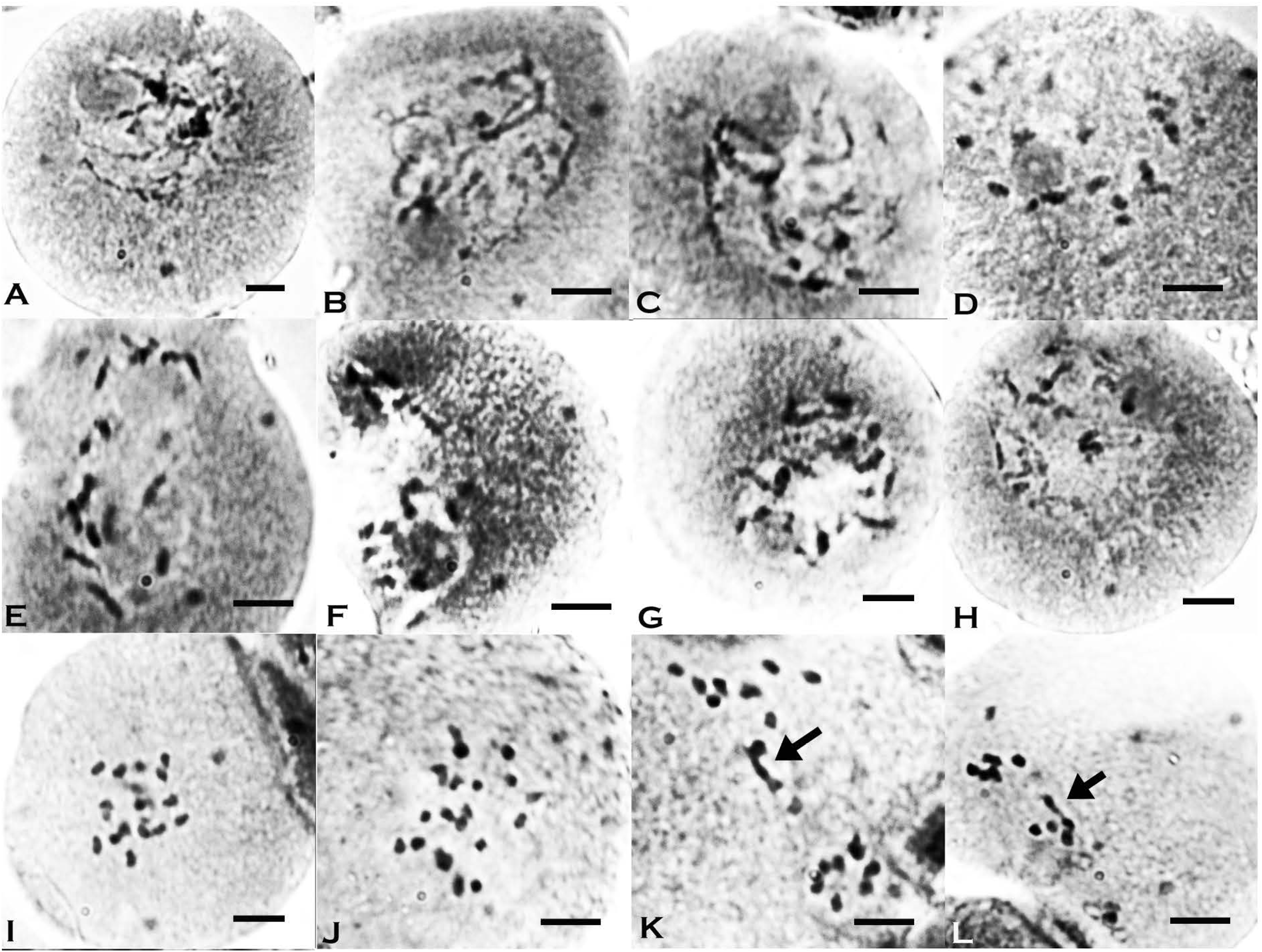
Acetocarmine staining of *spo11-1-1* meiocytes during meiosis I. **(A)** *spo11-1-1* meiocyte at pachytene. **(B-C)** *spo11-1-1* meiocytes at diplotene. **(D-H)** *spo11-1-1* meiocytes at diakinesis mostly exhibit univalents. **(I-L)** *spo11-1-1* meiocytes at metaphase I mostly exhibit univalents and occasional bivalents (arrows in K and L). Scale bar represents 10 μm.

**Figure S4.**
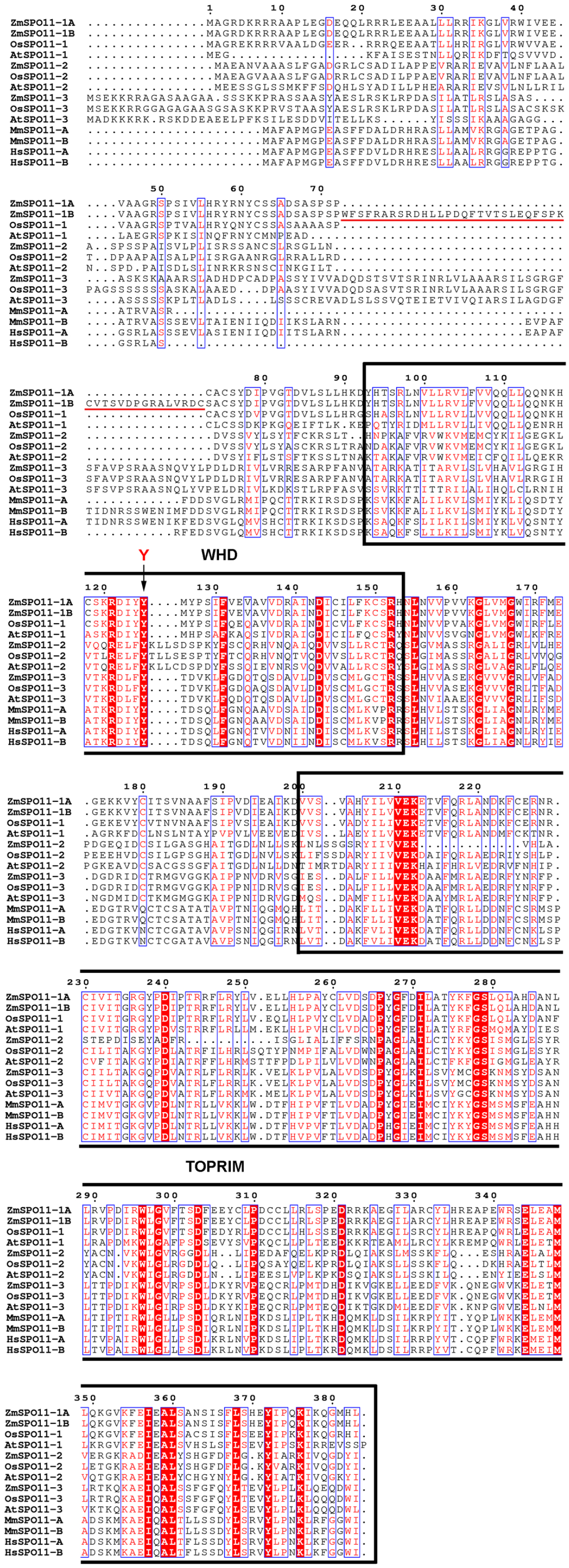
Alignment of SPO11 amino acid sequences from maize (Zm), rice (Os), Arabidopsis (At), mouse (Mm) and human (Hs). Conserved residues are highlighted in red. The conserved tyrosines (Y) in the WHD and TOPRIM domains are indicated. The additional 43-amino acid domain in SPO11-1β is underlined (red), which exhibits positional similarity to regions of SPO11-3 and the mammalian SPO11-β isoform.

**Figure S5.**
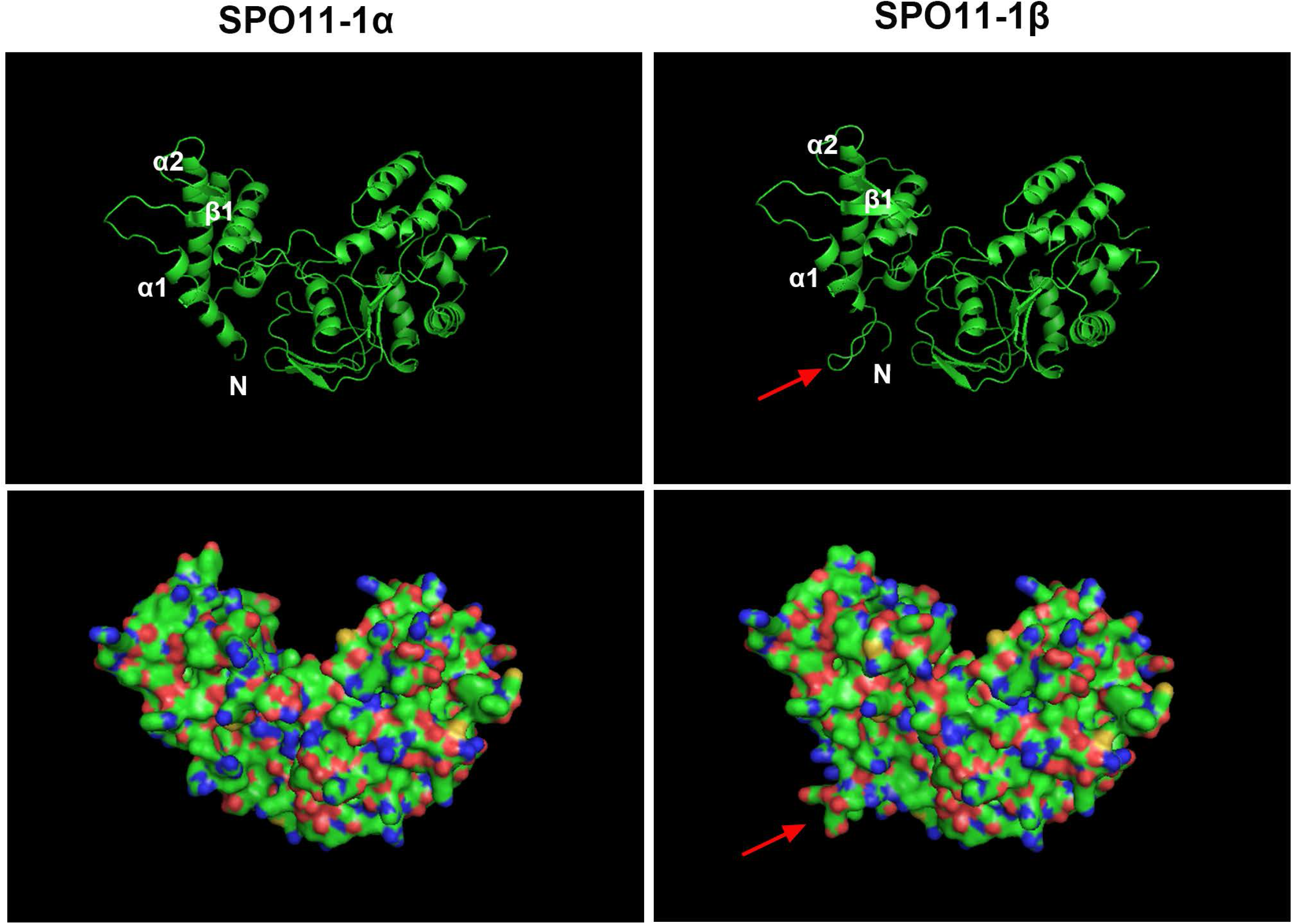
Predicted structures of the maize SPO11-1β and SPO11-1α isoforms. Predicted structures were obtained using Phyre2 and visualized using the PyMOL ‘cartoon’ (top) and ‘surface’ (bottom) tools. The SPO11-1 structure is based on the defined crystal structure of the TOPVIA protein of *Methanococcus jannaschii* (PDB model: c2zbkA). It forms a horseshoe shape that can dimerize into a ring. The additional domain of 43 amino acids in SPO11-1β manifests as a protruding alpha-helical region (arrows) opposite the groove containing the DNA binding region and the tyrosine catalytic site. N represents the N-terminal.

**Figure S6.**
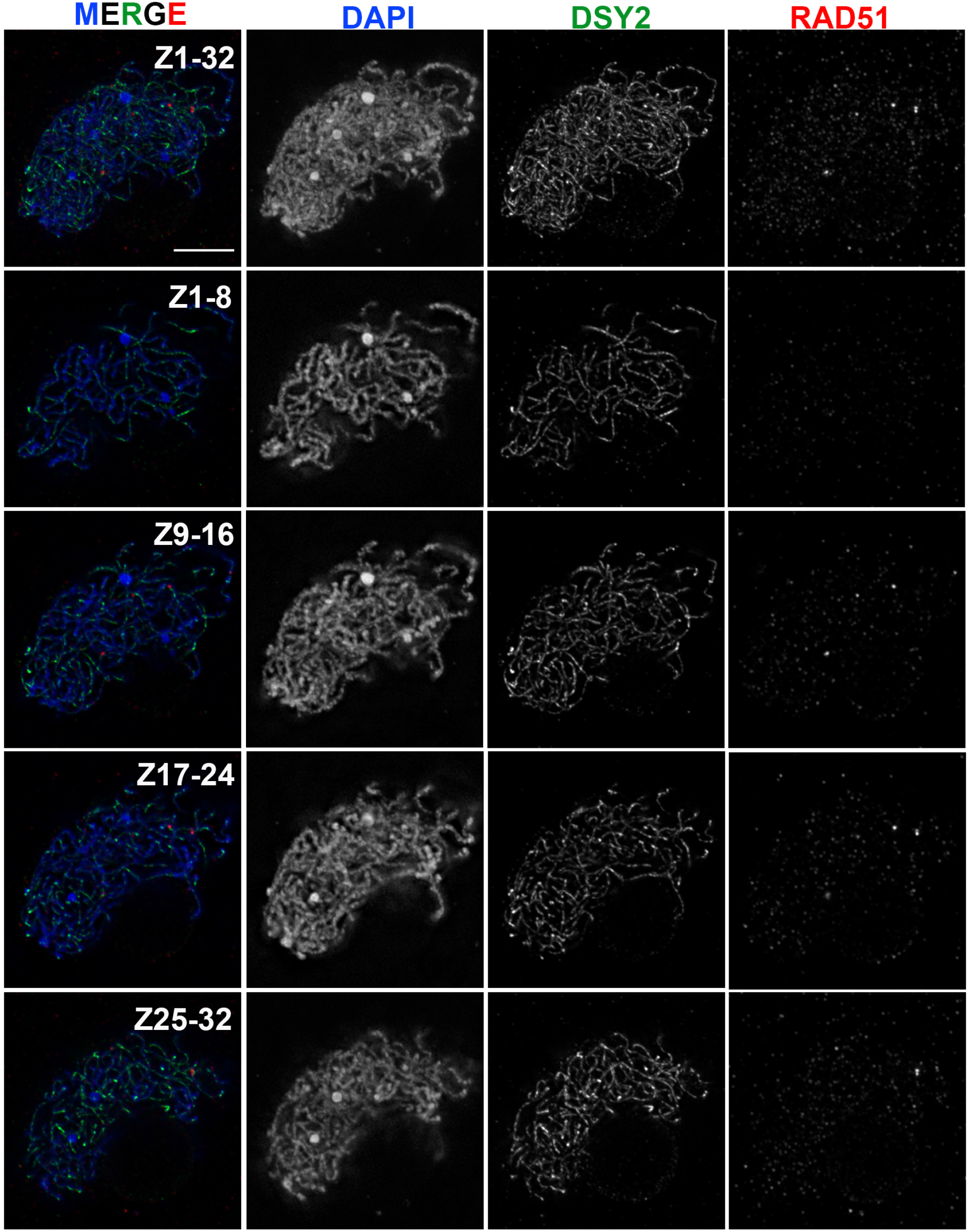
Approximately 10% of *spo11-1* meiocytes exhibit a few RAD51 foci. A representative nucleus of a *spo11-1-Δ03* meiocyte displaying a few RAD51 foci (red) is shown in projection images of the entire Z stack (Z1-32) and for different portions of Z stacks. Scale bar represents 5 μm.

**Figure S7.**
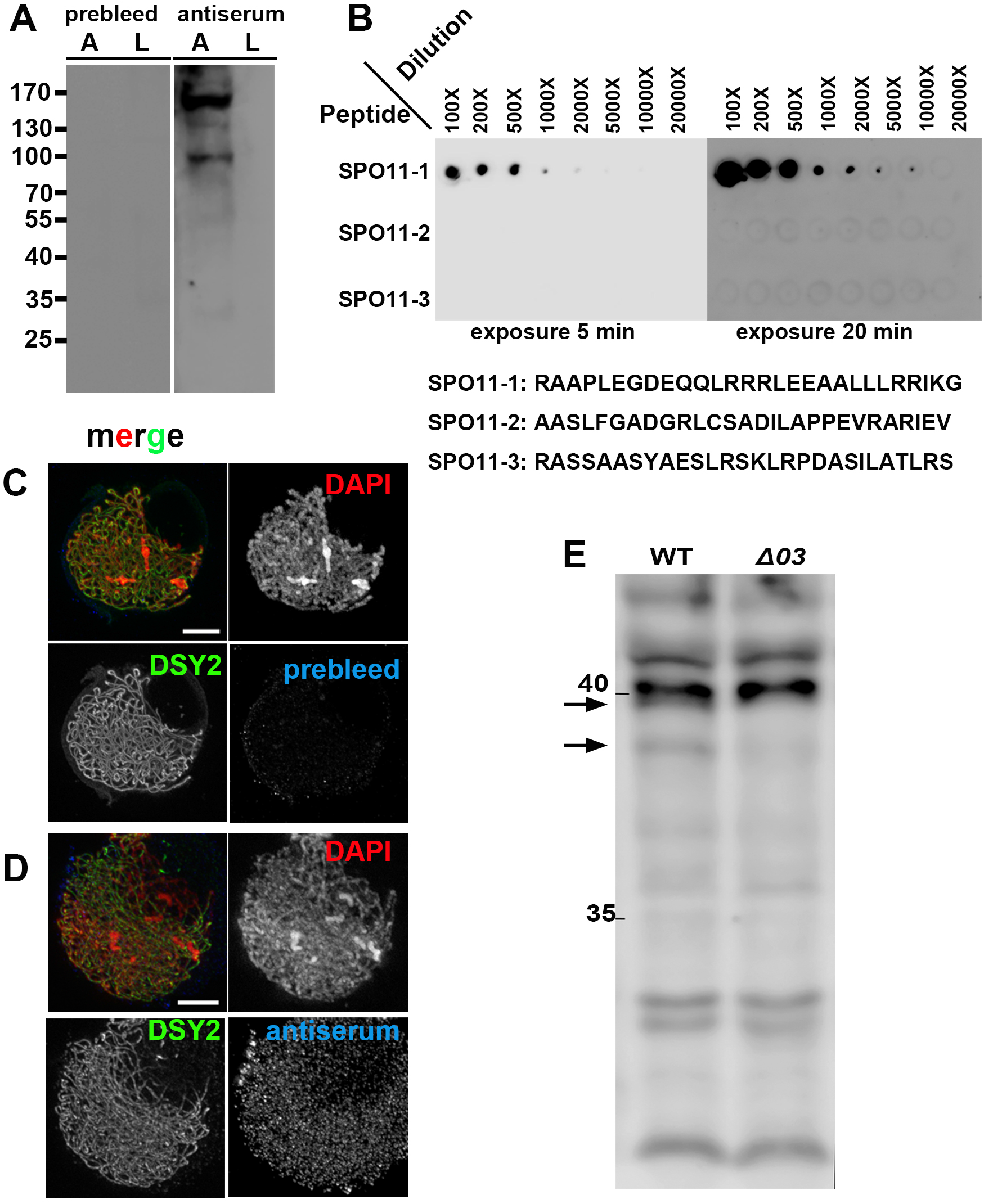
Generation of SPO11-1 antibody and validation of its specificity. **(A)** Western blot analyses with rabbit pre-immune serum (prebleed) and anti-immune serum (antiserum) were used to determine background levels before immunization and to detect any generated IgG that may recognize target proteins from total protein extracts of anther (A) and leaf (L) tissues. **(B)** Maize SPO11-1, SPO11-2 and SPO11-3 proteins share some similarities (Fig S4). To validate our SPO11-1 antibody specificity, dot blot analyses were performed using SPO11-1 antiserum (1:1000 dilution) against synthetic peptides of SPO11-1 antigen, SPO11-2 and SPO11-3 in corresponding regions. Their amino-acid sequences are listed below the dot blots. Serial dilutions of equal amounts (1 μg) of peptide were dotted for detection and blots were imaged using the UVP Biospectrum 600 system with exposure times of 5 or 20 min. **(C-D)** Pre-immune serum (C, prebleed) and anti-immune serum (D) were used to test SPO11-1 antibody in WT meiocytes at early zygotene stage by means of immunofluorescence analysis. **(E)** Western blot analysis using the affinity-purified SPO11-1 antibody for detection of potential SPO11-1 proteins in total protein extracts of WT and *spo11-1-Δ03* (*Δ03*) anthers. Two weak bands (arrows) of ∼40 kDa were detected in WT, but these were absent from the mutant sample.

**Figure S8.**
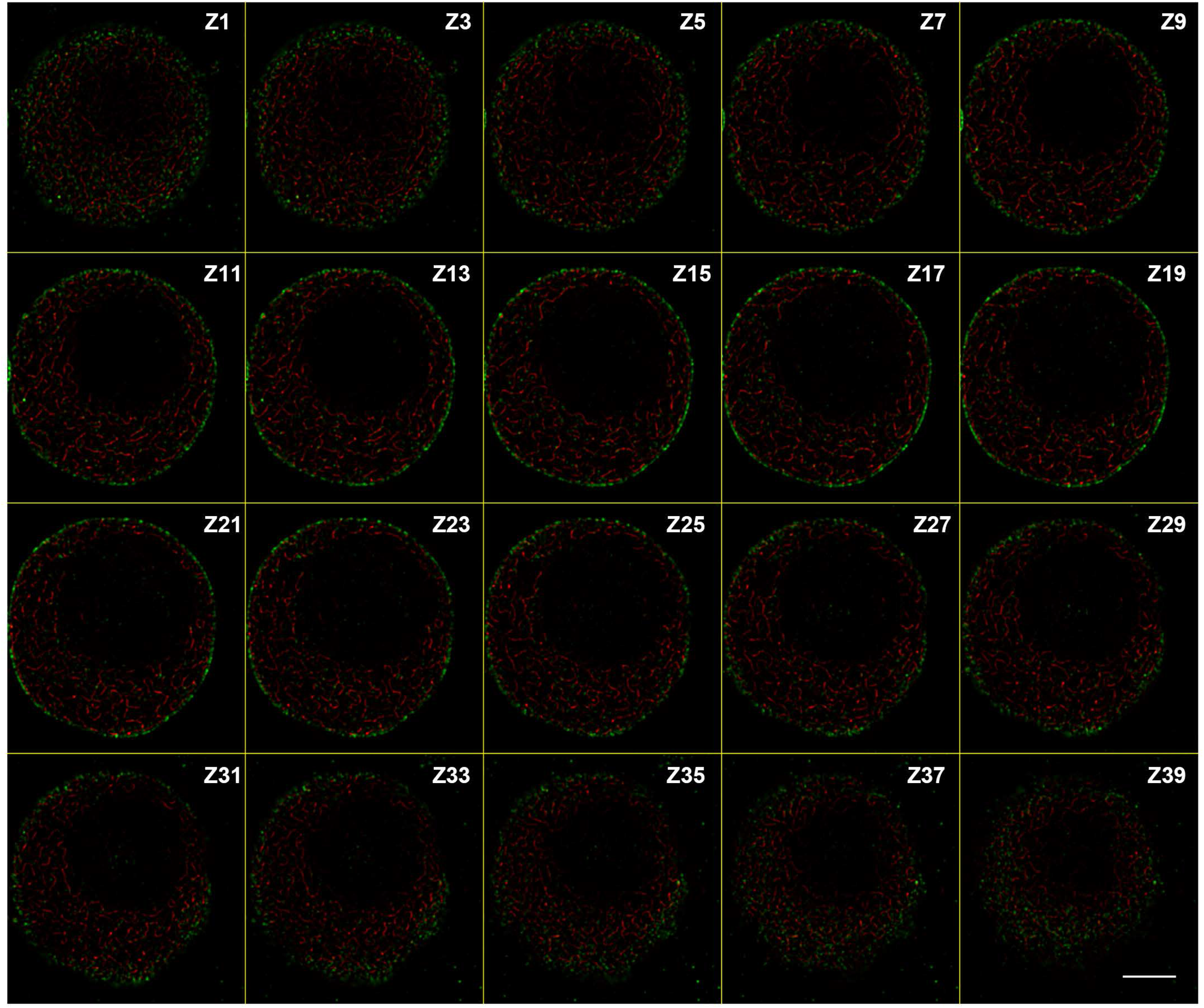
Montage of Z sections from the leptotene nucleus shown in Fig 5A. A montage of single Z sections with one Z-section intervals showing SPO11-1 (green) and DSY2 (red) signals. Scale bar represents 5 μm.

**Figure S9.**
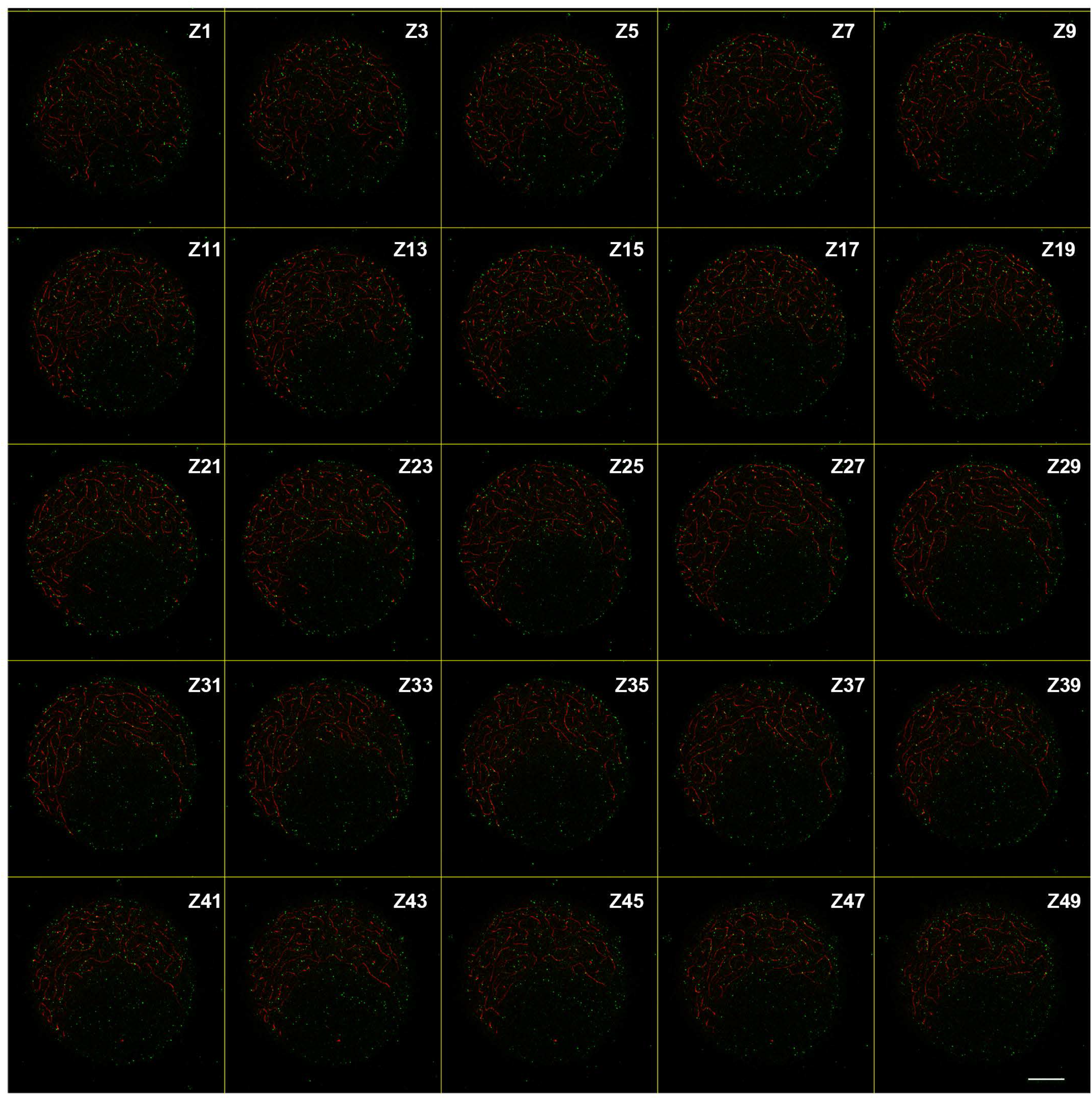
Montage of Z sections from the late leptotene nucleus shown in Fig 5B. A montage of single Z sections with one Z-section intervals showing SPO11-1 (green) and DSY2 (red) signals. Scale bar represents 5 μm.

**Figure S10.**
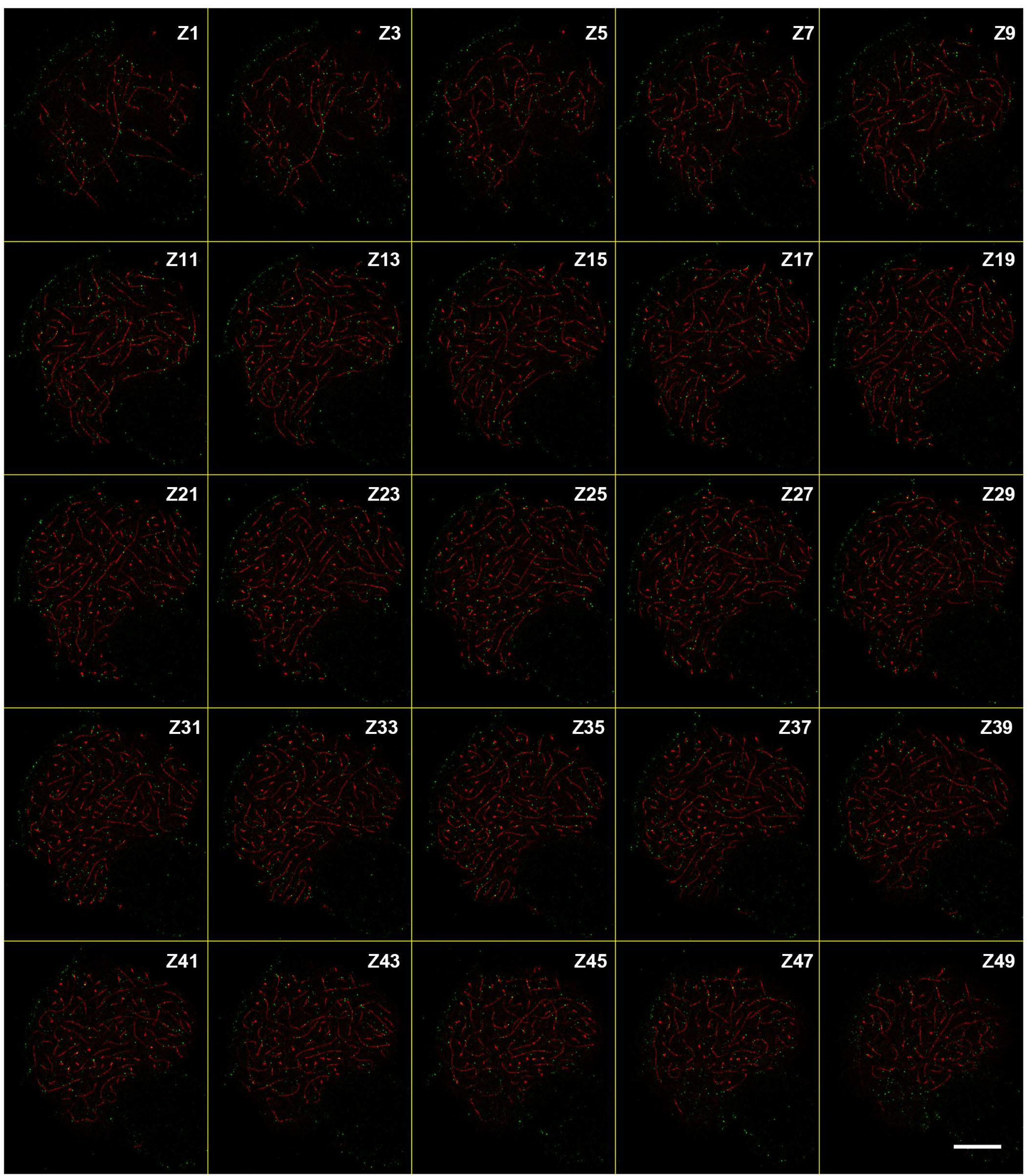
Montage of Z sections from the early zygotene nucleus shown in Fig 5C. A montage of single Z sections with one Z-section intervals showing SPO11-1 (green) and DSY2 (red) signals. Note that Z11-Z19 exhibit pre-aligned and synapsing regions. Scale bar represents 5 μm.

**Figure S11.**
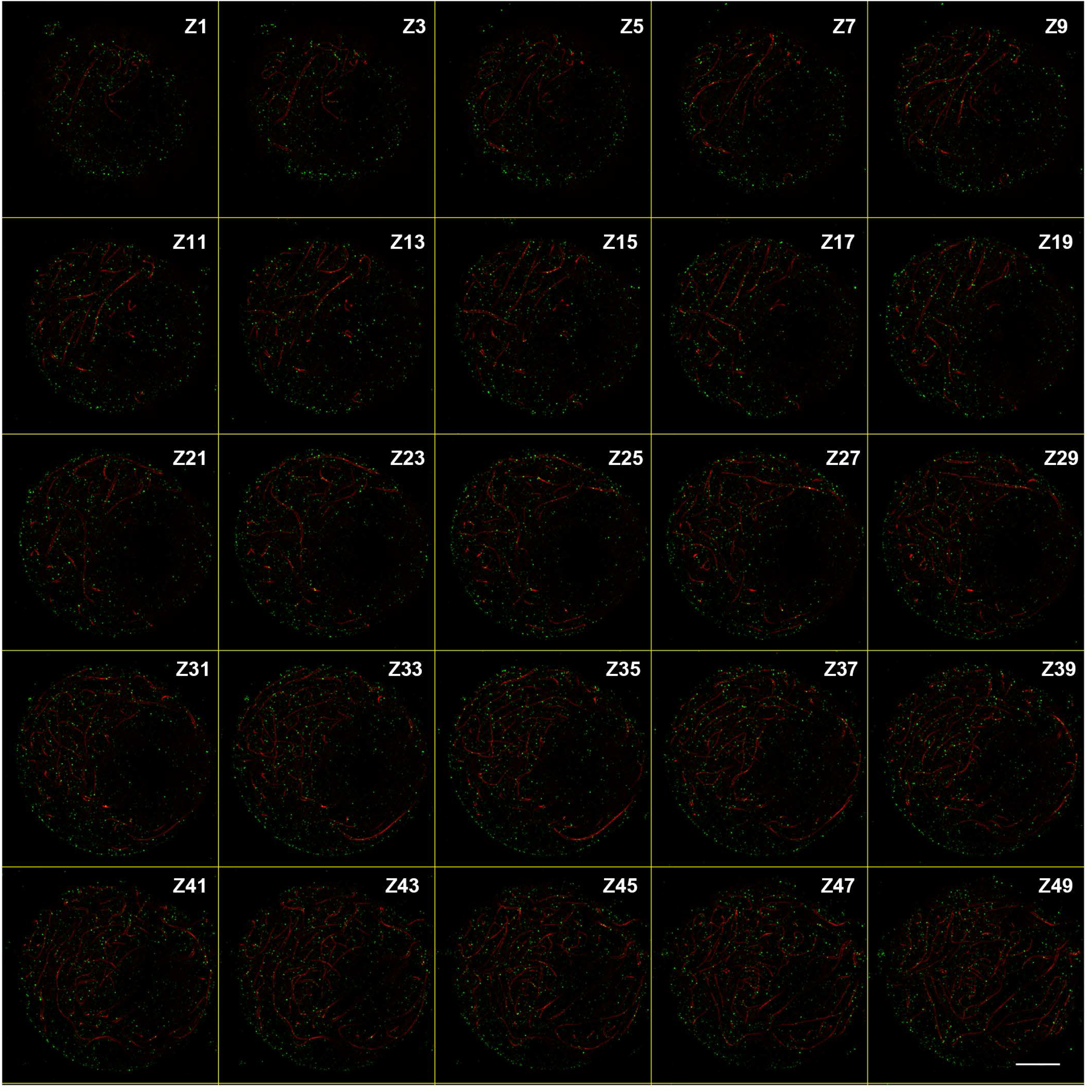
Montage of Z sections from the zygotene nucleus shown in Fig 5D. A montage of single Z sections with one Z-section intervals showing SPO11-1 (green) and DSY2 (red) signals. Scale bar represents 5 μm.

**Figure S12.**
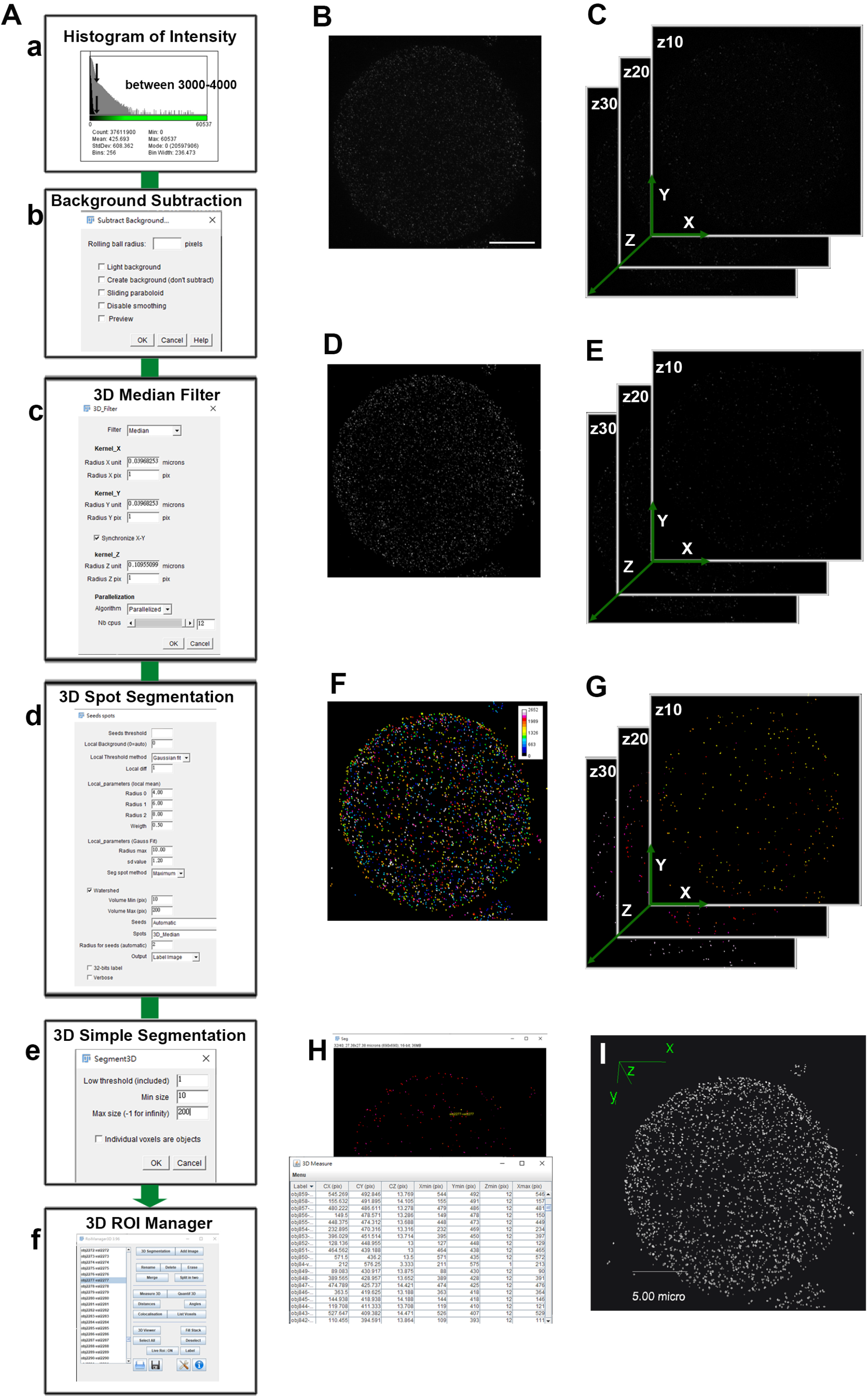
Overview of SPO11-1 signal segmentation. **(A)** Schematic workflow for segmentation and quantification of SPO11-1 signals. **(a)** Histogram of fluorescence intensity for all image stacks of a representative nucleus shown in B-C. The X-axis represents intensity values and the Y-axis shows the number of pixels found for each value. The black area displays a unimodal histogram, showing that the peak contains mostly lower intensity background, and the gray overlay shows a log-scaled version. The histogram of intensity is used to determine threshold values (arrows) that are usually between 3000-4000 for a 16-bit image. **(b-c)** Pre-processing step to enhance object contrast by rolling ball background subtraction and 3D median filtering. The resulting images are shown in D-E. **(d)** Detection and segmentation of SPO11-1 signals by 3D spot segmentation. using a threshold value defined by the histogram distribution in (a). This algorithm defines seeds of spots in original images using a threshold value defined by the histogram distribution in (a) and computed neighboring voxels as belonging to the object with criteria of volume of objects and local contrast (Gaussian fit). **(e-f)** Labeling of segmented objects by 3D simple segmentation and examination using 3D ROI manager. Geometric measurement of objects (H) was obtained using the Measure 3D tool. **(B-C)** Representative raw images of SPO11-1 signals shown in the maximal projection (B) and single Z sections (C). Scale bar represents 5 μm. **(D-E)** Pre-processed images of SPO11-1 signals shown in the maximal projection (D) and single Z sections (E). **(F-G)** Segmented objects shown in the maximal projection (F) and single Z sections (G). A total of 2652 3D objects was identified in the representative meiocyte. The color code represents numbered objects. **(H)** Labeled objects were inspected and measured using 3D ROI manager. **(I)** A surface-rendered image of SPO11-1 signals in the same nucleus visualized using 3D viewer.

**Figure S13.**
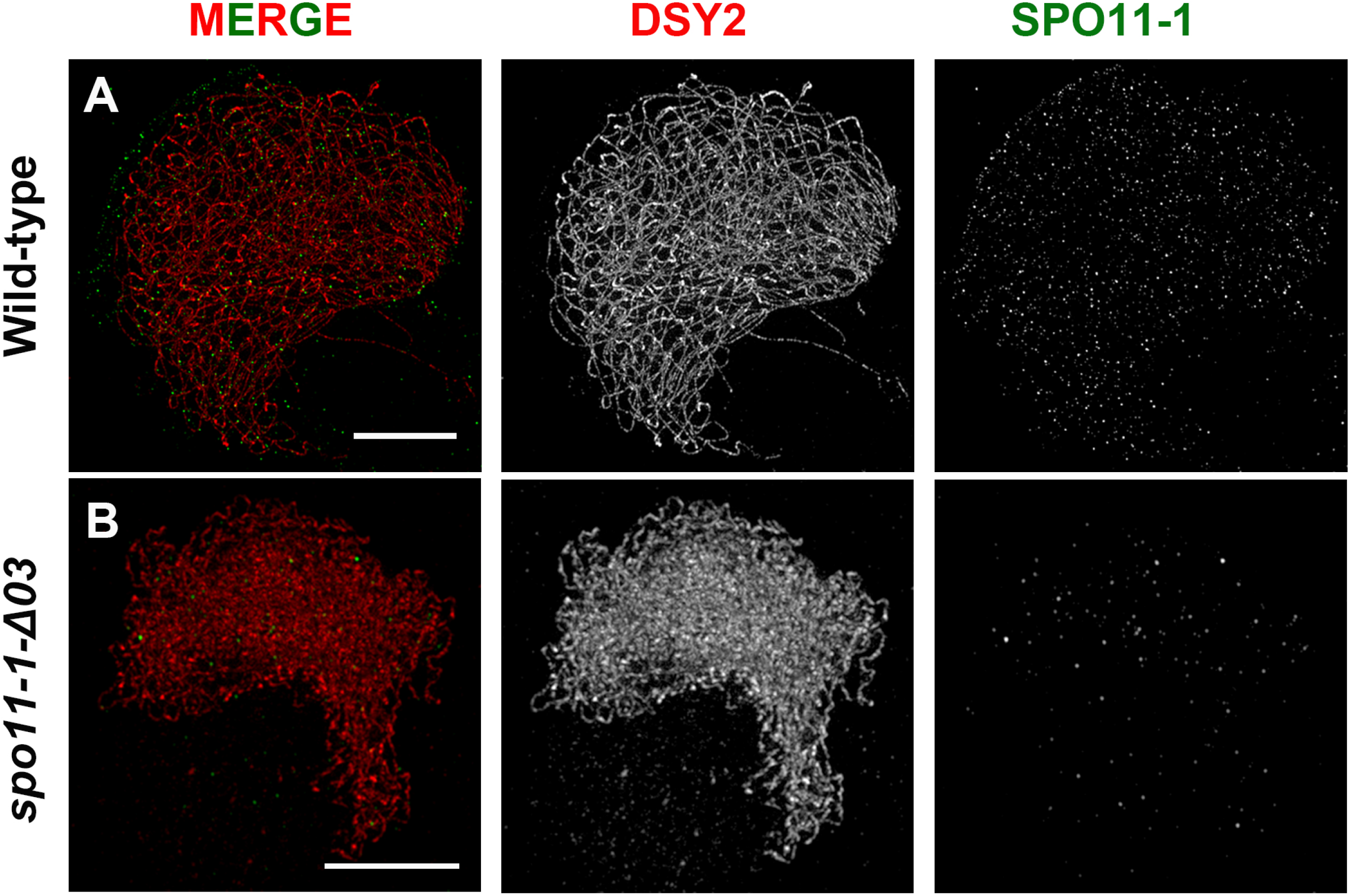
Super-resolution microscopy detects a few SPO11-1 foci in the *spo11-1-Δ03* mutant. **(A)** A WT meiocyte showing chromosome axes labeled by DSY2 (red or gray) and SPO11-1 signals (green or gray). **(B)** A representative *spo11-1-Δ03* meiocyte showing chromosome axes labeled by DSY2 (red or gray) and a few foci (green or gray) detected using a SPO11-1 antibody. Scale bar represents 5 μm.

**Figure S14.**
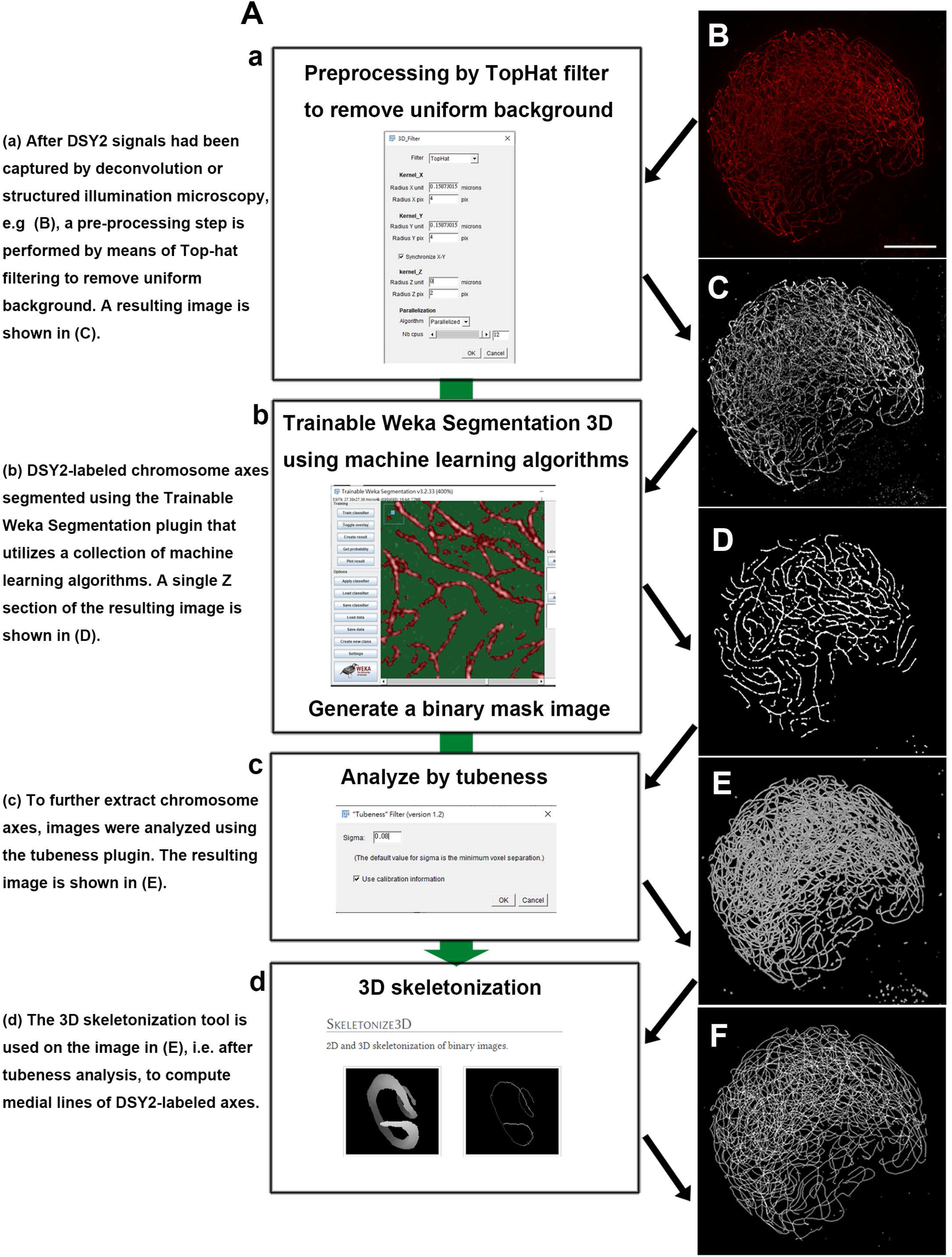
Segmentation of DSY2-labeled axial elements. **(A)** Schematic workflow for segmentation of DSY2 signals. **(a)** After DSY2 signals had been captured by deconvolution microscopy or structured illumination microscopy (B), a pre-processing step is performed by means of Top-hat filtering to remove uniform background. A resulting image is shown in (C). **(b)** DSY2-labeled chromosome axes segmented using the Trainable Weka Segmentation plugin that utilizes a collection of machine learning algorithms. A single Z section of the resulting image is shown in (D). **(c)** To further extract chromosome axes, images were analyzed using the tubeness plugin. The resulting image is shown in (E). **(d)** The 3D skeletonization tool is used on the image in (E), i.e. after tubeness analysis, to compute medial lines of DSY2-labeled axes. **(B)** A representative raw image of DSY2 signal in maximal projection. Scale bar represents 5 μm. **(C)** A pre-processed image of DSY2 signal in maximal projection after step (a). **(D)** Segmented axes from a single z section generated by step (b). **(E)** The resulting image of a representative nucleus in maximal projection after the tubeness analysis in step (c). **(F)** A skeletonized image of chromosome axes in maximal projection.

**Figure S15.**
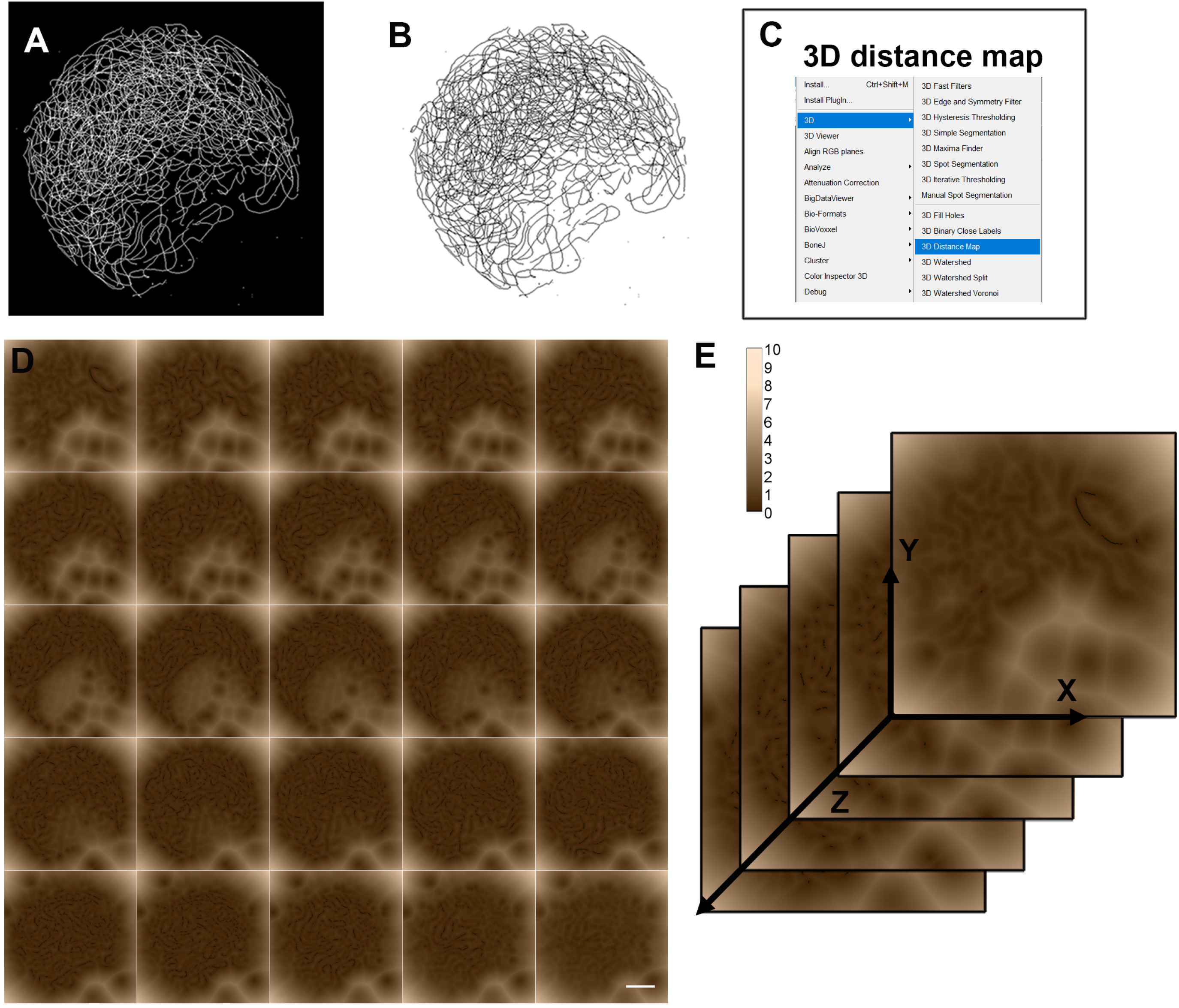
A 3D distance map computed from the skeletonized axis model. **(A)** A skeletonized image of chromosome axes in maximal projection. **(B)** The same image (A) converted to white background. **(C)** Screenshot of the 3D distance map tool in ImageJ. **(D, E)** A 3D distance map was generated from the skeletonized axis model (B) using the 3D distance map tool. These maps could be visualized in 2D by using color to denote the distance from a given point (i.e. medial lines of the skeletonized axis) to neighboring voxels. A montage of serial z sections (D) and a schematic image of Z stacks (E) of the resulting 3D distance map are shown. The color scale represents assigned distances (μm).

**Figure S16.**
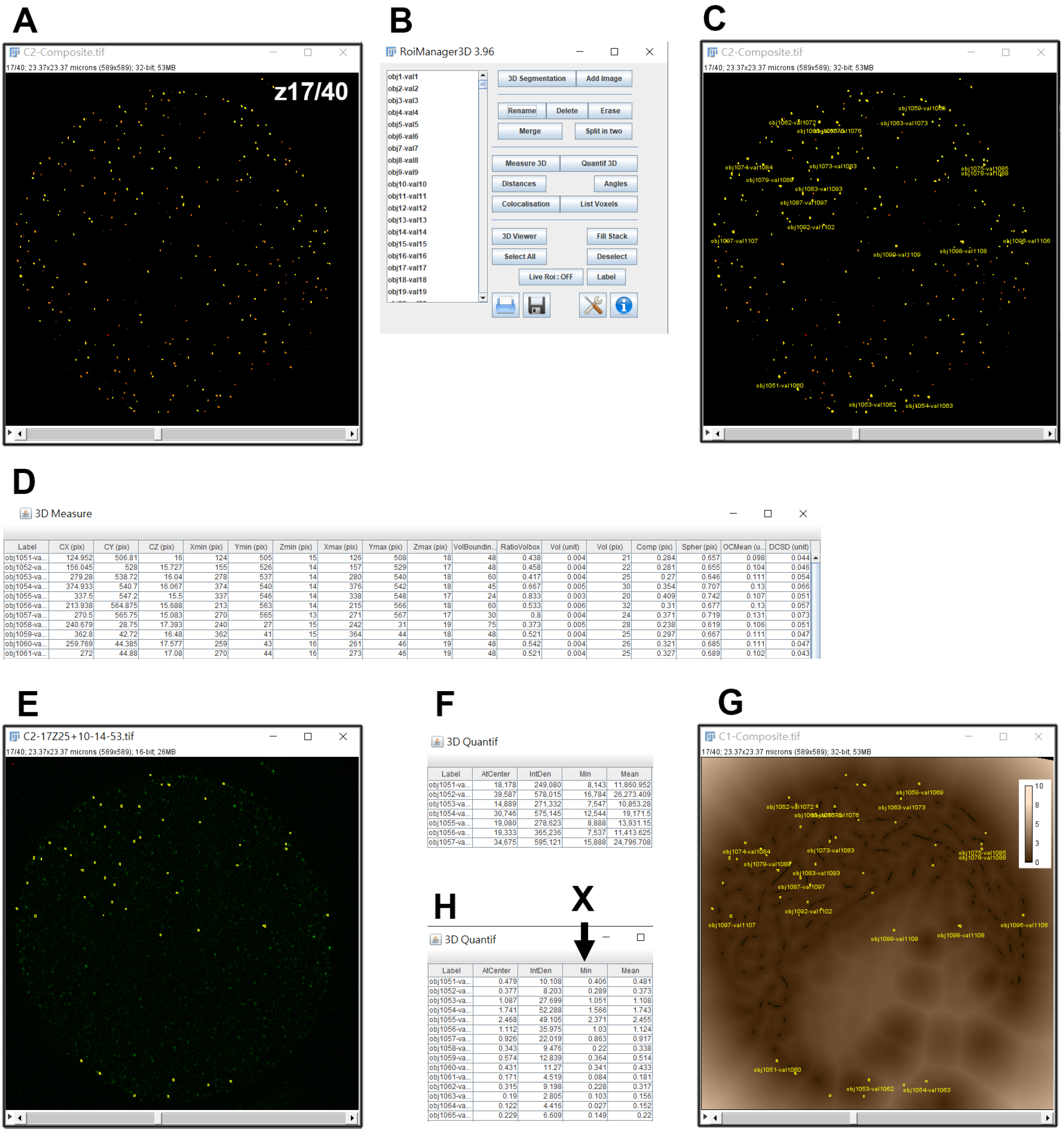
Calculation and measurement of distances between SPO11-1 and skeletonized axis. **(A)** A representative image of segmented SPO11-1 signals shown in a single Z section. The segmentation method for the same cell is demonstrated in Fig S12. **(B)** The 3D ROI manager tool is opened and the segmented binary image is imported to add all labeled objects. **(C)** Examination of selected objects in the segmented binary image using 3D ROI manager. **(D)** Acquisition of geometric measurements of selected objects, including coordinates of the center of objects (CX, CY, CZ), the borders of objects, their volumes (Vol) and compactness (Comp), as well as the distances from the centers to borders of objects (DC). **(E)** Examination of selected objects (circled in yellow) from the original gray-scale image. **(F)** Acquisition of the fluorescence intensity of selected objects from the original gray-scale image. Signal intensity values for the center point (AtCenter), total sum (IntDen), minimal (Min) and the mean (Mean) for the objects are obtained. **(G)** After incorporating the 3D distance map into the 3D ROI manager, selected objects (circled in yellow) could be examined on the 3D distance map. The segmentation and skeletonization method for chromosome axes of the same cell is demonstrated in Fig S14. **(H)** Acquisition of distances (μm) between the centers of selected objects to the closest axes (AtCenter), as well as minimal (Min) and mean (Mean) distances from the borders of objects to the closest axes. Axis-associated SPO11-1 objects are defined when X values (i.e. the minimal distance from AEs to borders of an object) equal zero.

**Figure S17.**
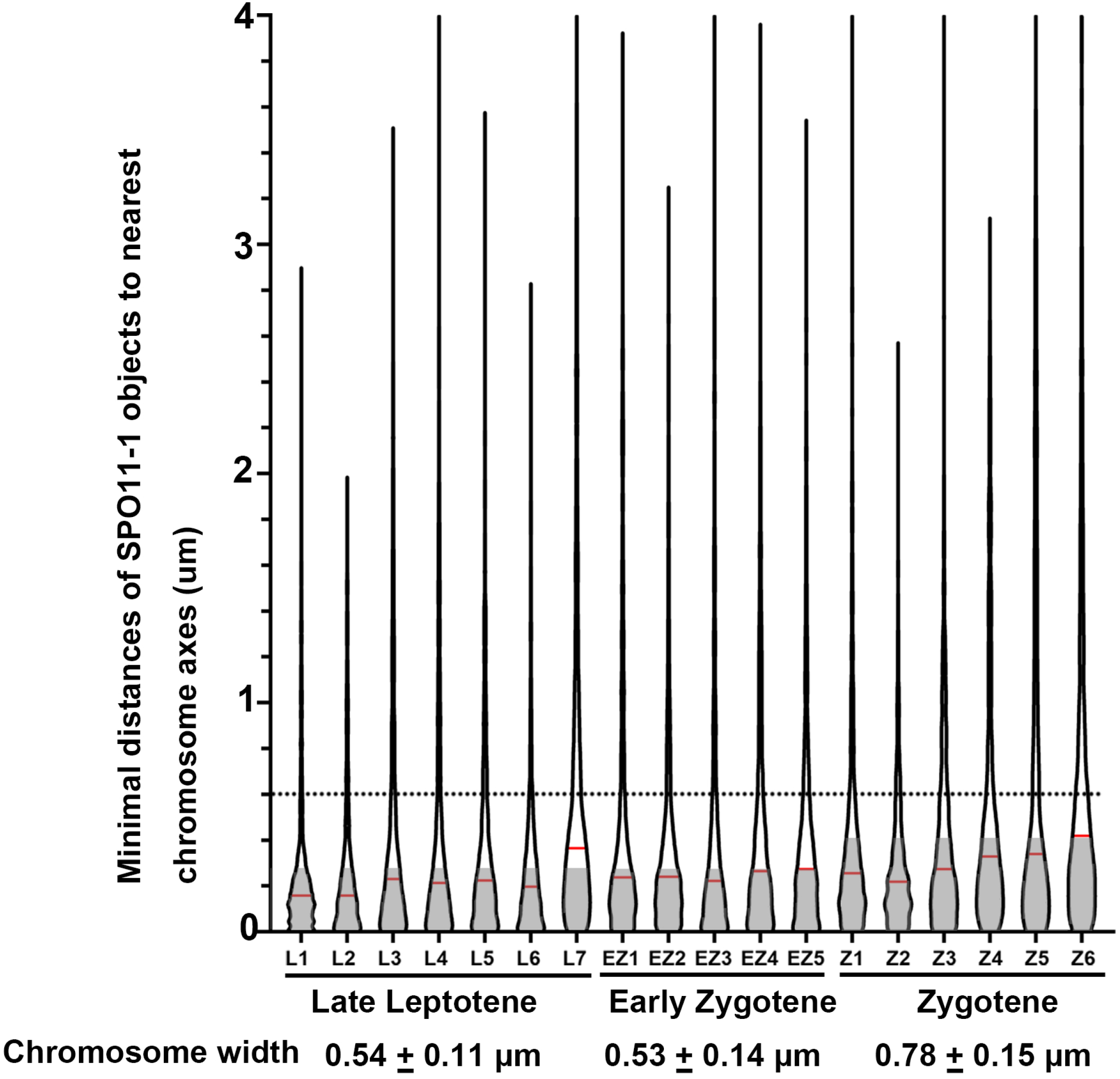
Distribution of distances (μm) between each SPO11-1 object to the nearest axis of a meiocyte. A total of eighteen meiocytes was analyzed, including seven late leptotene (L), five early zygotene (EZ) and six zygotene (Z) cells. Median values are indicated by red lines. The dotted line represents a radius of 0.6 μm from an axis. The shaded area represents estimated DAPI-stained chromosomal regions away from the axes (i.e. half the value of chromosome width). Chromosome widths were estimated by measuring at least 100 positions.

**Figure S18.**
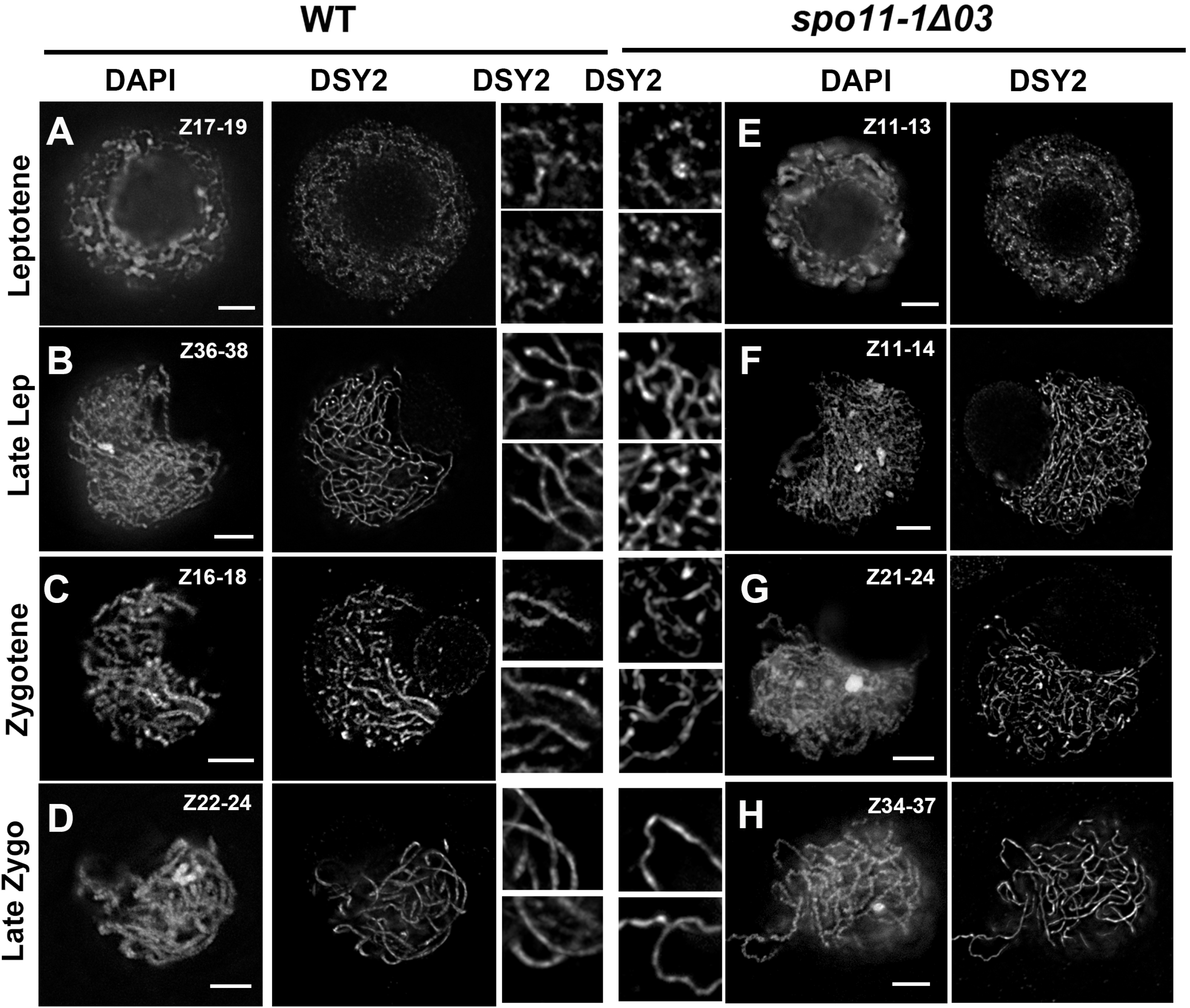
Immunolocalization of the axial element protein DSY2 in wild-type and *spo11-1* mutant meiocytes. **(A-D)** Maximal projection images of a few Z sections from WT meiocytes during early prophase I. DAPI staining and DSY2 signals are presented separately. Scale bar represents 5 μm. Two magnified regions (5 μm) of DSY2-labeled chromosome axes are shown in the panels at right. **(E-H)** Maximal projection images of a few Z sections from *spo11-1-Δ03* meiocytes during early prophase I. DAPI staining and DSY2 signals are presented separately. Scale bar represents 5 μm. Two magnified regions (5 μm) of DSY2-labeled chromosome axes are shown in the panel at left.

**Figure S19.**
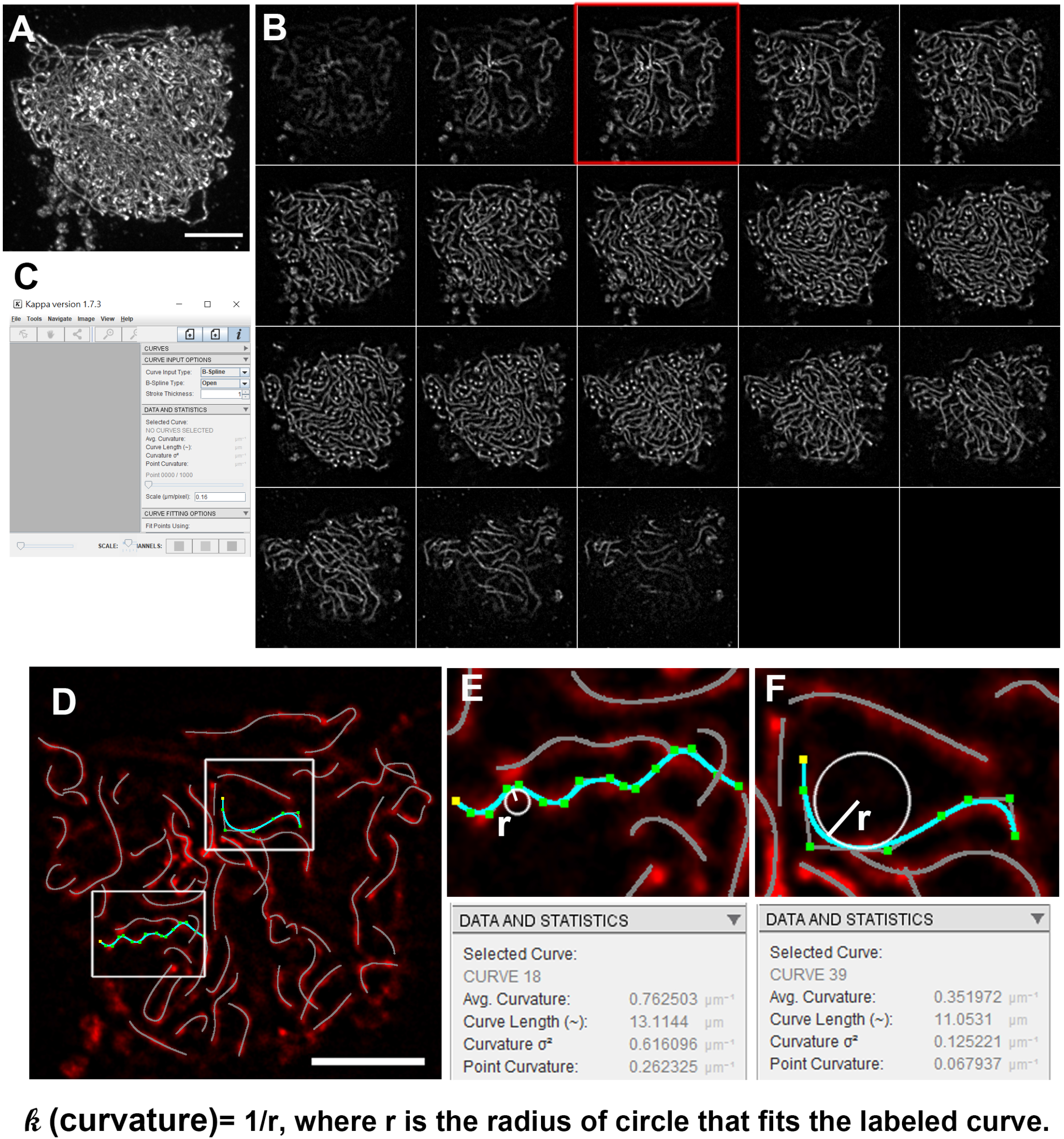
Measurement of axial element curvatures. **(A)** Immunostaining image of DSY2 signal in maximal projection. Scale bar represents 5 μm. **(B)** A montage of single Z sections. **(C)** The Kappa plugin in ImageJ. **(D)** An example of one Z section (indicated by the red box in B) subjected to Kappa analysis. DSY2 stretches are traced and marked by blue lines. **(E and F)** Magnified regions in D. Measurements of the labeled curves are shown below the screenshots. The Kappa value of the curve is calculated by 1/r, where r is the radius of a circle that fits the points along the curve. A bigger Kappa represents a curve bend more sharply. As shown here, average kappa values of curve 18 (E) and curve 39 (F) are 0.762503 and 0.351972 μm^-1^, respectively.

**Figure S20.**
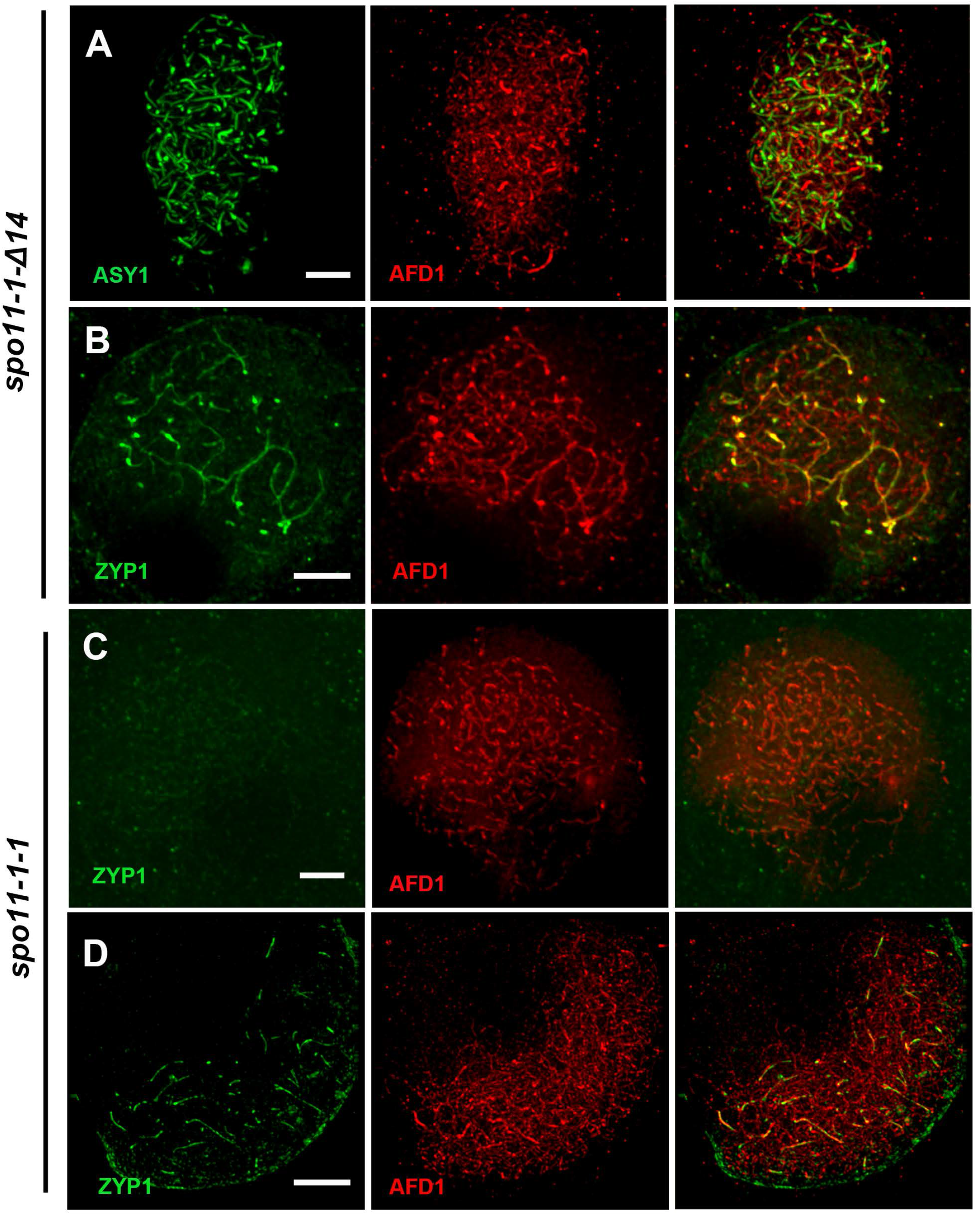
Immunolocalization results in *spo11-1-Δ14* and *spo11-1-1* meiocytes reveal similarly abnormal phenotypes. (A, B) Detection of cohesion protein AFD1/REC8, axial element ASY1 and transverse filament ZYP1 proteins at early-zygotene (A) and late-zygotene (B) of *spo11-1-Δ14* mutant meiocytes showing aberrant AEs (A) and impaired SC formation (B). (C, D) Detection of cohesion protein AFD1/REC8 and transverse filament ZYP1 proteins at zygotene (C) and late-zygotene (D) in *spo11-1-1* mutant meiocytes showing impaired SC formation. Scale bar represents 5 μm.

**Figure S21.**
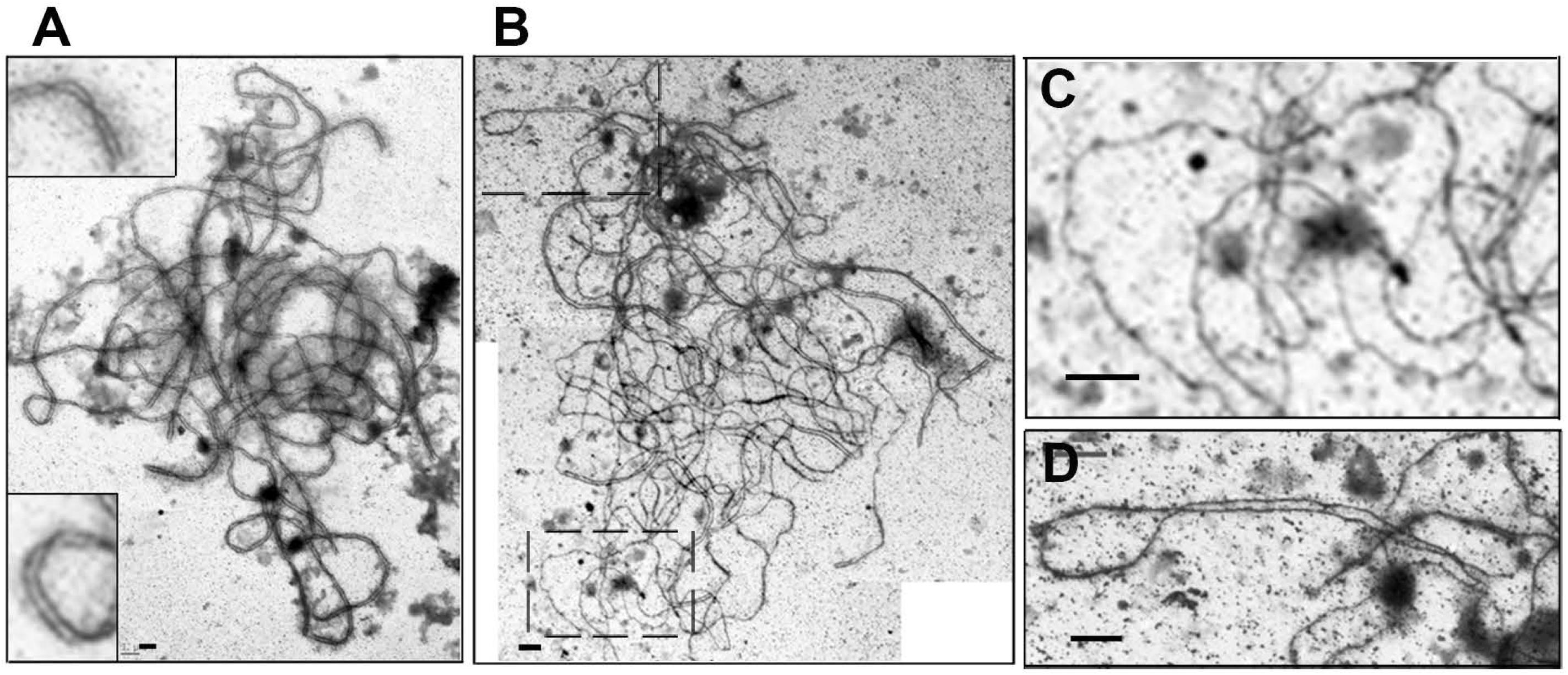
Transmission electron microscopy of synaptonemal complex chromosome spreads of *spo11-1-14Δ* mutants. **(A)** WT pachytene cell exhibiting complete synapsis. **(B)** Promiscuous synapsis in the *spo11-1-Δ14* mutant. A stitched image is shown here due to the oversized spread of the synaptonemal complex of maize meiocytes. **(C, D)** Higher magnifications from (B). Abnormally curly AE in unsynapsed regions (C). Abnormal fold-back synapsis (D). Scale bars represent 1 μm.

